# The carbon footprint of meat and dairy proteins: a practical perspective to guide low carbon footprint dietary choices

**DOI:** 10.1101/2021.01.31.429047

**Authors:** R. Gaillac, S. Marbach

**Author notes:** Electronic mail (S. Marbach).

## Abstract

Meat and dairy products in the food industry represent a significant portion of anthropogenic green house gas emissions. To meet the Intergovernemental Panel on Climate Change recommendations to limit global warming, these emissions should be reduced. Meat and dairy products are also responsible for the majority of our daily, vital, protein intake. Yet, meat and dairy products contain very different amounts of proteins, making it difficult in general to rationalize which protein source has the lowest carbon footprint. Here we present a practical and pedagogical review, comparing the carbon footprint of a variety of meat and dairy products with respect to their protein content. We investigate the carbon footprint of different dietary choices for several countries, by keeping the total number of meat and dairy proteins constant. Interestingly, we find that dairy-only diets are in general only a little less carbon intensive than current diets. However, 50% carbon footprint reduction may be obtained, throughout the world, with a “low CO_2_”-tailored diet including only small poultry, eggs and yogurt. Such a dietary pattern suggests easy to follow consumer guidelines for reduced carbon footprint. We report further on a number of consumer oriented questions (local or imported? organic or not? cow or goat milk? hard or soft cheese?). Our methodology may be applied to broader questions, such as the carbon footprint of proteins in general (including fish and plant proteins). We hope our work will drive more studies focusing on consumer-oriented questions.

## I. INTRODUCTION

Climate change, resulting from the emission of greenhouse gases by human activities – in particular carbon dioxide, is a worldwide threat with long-lasting implications^1^. To limit the increase of global average temperature compared to pre-industrial level, substantial efforts have to be made. Indeed, according to the Intergovernemental panel on climate change (IPCC), limiting global warming to 1.5°C requires to reduce the emissions by 45% from 2010 levels by 2030, and to reach net zero by 2050^1^ – see Fig. 1. Even limiting global warming to 2.0°C brings these numbers to a 25% decrease of emissions in 2030, and to reach net zero in 2070^1^.

**FIG. 1.**
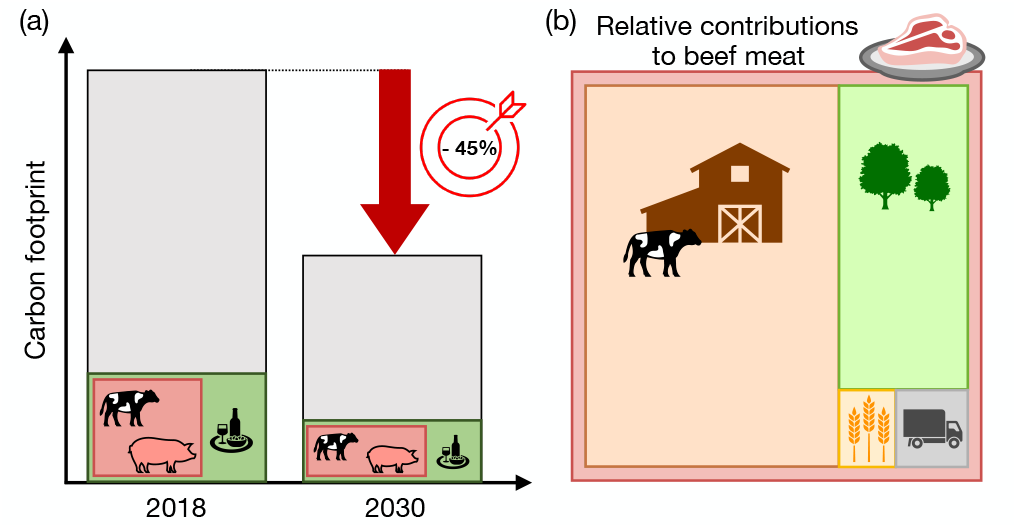
(a) Proportion of food and livestock based products in the global carbon footprint balance of anthropogenic emissions and the relative change recommended by IPCC for the 2030 Target. The areas of the respective boxes correspond to the proportional carbon footprints of food production and livestock management. (b) Example of relative carbon footprint contributions for the production of beef meat at the farm (orange), land use change (green), feed production (yellow) and transport (gray). Similarly, box areas correspond to relative carbon footprint contributions.

### A. A brief on CO_2_ emissions of food

Per year, the food supply chain generates 13.7 billion metric tons of carbon dioxide equivalents (CO_2_ eq.)^2^. That represents 26% of the total anthropogenic green-house gas (GHG) emissions^2^ – see Fig. 1-a. Furthermore, significant increase of food chain related emissions is expected with population increase and income level increase^3^. Therefore, in line with IPCC guidelines, reducing the emissions of the food supply chain is critical^3–5^.

Among the food supply chain, meat and dairy production generates a significant amount of GHG emissions. Livestock alone represents at least 14% of the total world emissions^5–7^. More than half of the emissions from food stems from livestock because a number of production steps are carbon intensive. For example, to produce beef, everything that happens at the farm (methane emissions from cows, farm machinery) represents on its own 66% of the emissions^2^. Land use change (initial deforestation to create a pasture, and subsequent soil contamination) represents 27 %^2^ and animal feeding (growing crops to feed livestock) represents 3 %. Transport, processing, packaging and retail fill up the remaining categories (so the remaining 4%) – see Fig. 1-b.

But just how much meat does that represent in the consumer’s plate? Meat consumption for an American averages to 120 kg of meat per year^8^, corresponding to about 340 g of meat per day (not counting food losses at the consumer lever)^1^. This value falls to 210 g/day^8^ in the European Union, 160 g/day in China and the world average is 115 g/day^8^. Calorie wise, taking a typical number of 200 kcal*/*100g of meat^10^, beef represents 8-24% of total calorie intake^11^^2^. This is a relatively small fraction considering it accounts for more than half of the carbon footprint. This imbalance between actual calories provided and carbon footprint can be further illustrated by the following number. In the United States (US), 4% of food sold (by weight) is beef, but that represents 36% of food-related emissions in the country^12^.

All in all, meat and dairy products represent the most relevant food category contributing to the total carbon footprint of dietary choices. A critical common point of meat and dairy products is that they are foods with high protein content, and are therefore primary sources of protein in current diets. In the following, note that we will also include eggs in the “dairy” category as they represent a significant source of protein in common diets.

### B. What are proteins, why do we need them and just how much ?

Proteins are large molecules made up of chains of amino acids. When we digest proteins, we break them down into amino acids – see Fig. 2. Amino acids achieve vital functions in our body – for example some are used for neurotransmission^13^. Amino acids can be further broken down to produce energy to power our body^14^ (and the rest of the pieces – urea and carbon dioxide – are eliminated by urine and breathing). Finally they can also be reassembled by the organism to synthesise other kinds of proteins that achieve a number of other vital functions in our body^13^. In short, it is impossible to live without proteins.

**FIG. 2.**
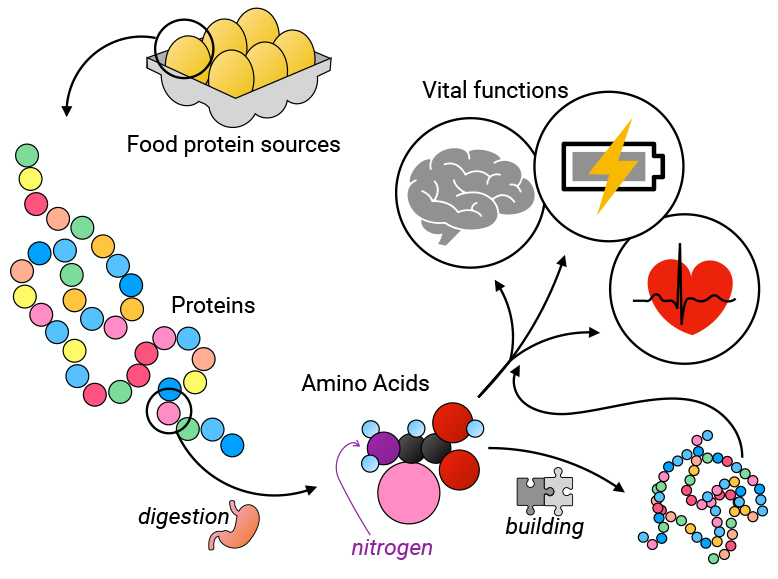
Illustration of the cycle of proteins and amino acids in human nutrition and their use in several vital functions.

Typically, for a person in good health (and that does not do any major sport training), the globally established dietary reference intake is about 0.8 g of protein per kilogram of body weight per day^11,15^)^3^. This means that a person weighing 60 kg (132 pounds) needs about 48 g of proteins per day, or a person weighing 80 kg (176 pounds) needs 64 g of proteins per day.

Higher values of protein intake per day can be beneficial in some circumstances. Up to 2.0 g*/*kg*/*day is beneficial to maximise muscle protein synthesis in resistance-training adults, with a maximum of 0.4 g*/*kg*/*meal^17^. Furthermore it is a common misbelief that a high protein diet – alone – can impact bone health^18, 4^. For the elderly, muscle strength preservation can be improved by protein intakes up to 1.0 g*/*kg*/*day accompanied by safe endurance and resistance type exercises^19,20^.

High *animal* protein intakes may however be connected with some specific diseases. For instance, high intake of *animal* protein – from 0.8 g*/*kg*/*day and over – may be connected to some age-related diseases (cancer, diabetes, cardiovascular diseases)^21–24^. This is especially true for red meat (beef, pork, mutton and lamb) and even more so for processed meat^21,25^. Substitution of animal protein by plant protein is beneficial to reduce overall mortality^23^. Of course, this substitution must still result in an adequate protein intake. Indeed an insufficient protein intake can also yield age-related diseases, especially muscle loss^21^. Note that adequate protein intake from plant sources is possible as all necessary amino acids may be found in plant based foods (especially in soy and legumes like lentils)^15,26^^5^.

To put these numbers in perspective, in the US, it appears in average people eat 1 −1.5 g*/*kg*/*day of protein^27^^6^. Out of these, about 60% are meat and dairy sourced proteins^29^, and therefore meat and dairy represent the most important source of proteins in current diets. Thus it is only natural to investigate the carbon footprint of meat and dairy proteins.

### C. Scope of this study: consumer oriented review to guide low carbon footprint choices among meat and dairy proteins

For all these reasons, we investigate here the carbon footprint of meat and dairy proteins. Inspired by the works of Ref. 2, 30, and 31, we aim for a measure of the carbon footprint per gram of protein for these different sources. We expose our methods in Sec. II. This allows us to directly compare different sources of proteins (among meat and the variety of dairy products) and determine which ones have the lowest carbon footprint (Sec. III). This is especially relevant as it directly answers the consumer question of which product to select for climate change mitigation, as far as proteins are concerned. Note that this is the first analysis that directly compares the carbon footprint of protein rich foods yet with very different protein contents (such as milk with 3, eggs with 10 and meat with 20 g protein/100 g edible food)). We also investigate a very broad range of products, in particular through the comparison of milks from different ruminants (cows, goats, sheeps, and buffalo) and the variety of cheeses. This is in sharp contrast with former studies that are focused on meat^2,30,31^.

We take our analysis closer to consumers by investigating specific dietary changes to reduce carbon footprint, taking into account regional discrepancies both in dietary patterns and in carbon footprints of products (Sec. IV). This allows us to find a “low CO_2_” diet guide, that consists only of small poultry (chicken, duck, rabbit), eggs and yogurts. This “low CO_2_” diet, while providing the same total amount of proteins than reference diets, enables to reduce carbon footprints by 50%, reaching the IPCC 2030 target. Such a diet is a reliable guide notably across the world. Importantly, we have identified that vegetarian diets (with high amounts of dairy proteins) are not nearly as effective (only 20% reduction) as our “low CO_2_” alternative. This highlights that as protein sources, dairy products in general do not have a low carbon footprint.

Finally, we discuss a number of consumer-type questions associated with meat and dairy consumption: such as the choice between local or imported products, organic or non-organic, nutritional questions and methodological questions (Sec. V).

We stress again that here we focus specifically on meat and dairy proteins. As mentioned earlier, not only do they represent the most abundant source of protein and the part of our diets with the largest carbon footprint, but a number of relevant consumer-oriented questions have to be addressed for these food categories. Fish and plant proteins are beyond the scope of this review. Finally, in line with our desire to answer consumer-oriented questions, we have adopted a pedagogical style throughout.

## II. METHODS

For this study we retain only the most common meat and dairy protein-rich products (discarding especially those for which data availability is limited). Among meat products, we explore the carbon footprint of beef, lamb, veal, pork, turkey, chicken, rabbit, duck and among dairy products we explore cow-based dairy: milk, cheese and yogurt; and finally chicken eggs. Different dairy sources (such as goat, sheep and buffalo) and the variability of dairy products (different cheeses and yogurts) are also compared.

### A. World wide data

#### 1. Protein content

Protein content ranges for the products investigated were taken from various national databases^10,32–34^ – making sure that the methods for protein quantification in foods^35^ were consistent.

Protein content data has a lot of variability. For example, the breeding methods used change with time and affect the protein content^32^. But also the breed itself and the sex of the animal^36^. Moreover, ready-to-eat meat comes from different parts of the animal that do not have the same content in water and fat and therefore the content in protein differs (such as sausage for which the fat content is higher in average, and therefore less dense in protein than trimmed steak). Finally, extrinsic properties caused by manufacturing and processing affect the protein content^32,36^. All of these factors also affect what is said to be the “meat quality”. Meat quality is a measure of the different kinds of amino acids (coming from proteins) that can be found in the meat and how they are ingested and properly used by our organism^10,26,36^. To lessen such variability, here we discard processed foods such as patties, sausages, and other prepared meals.

This allows to retain a range of protein content *P*_min_ − *P*_max_ for each product.

More information on the methods and the data retained in this study can be found in Appendix A.

#### 2. Carbon footprint

Worldwide data on carbon footprint of the products investigated has been analyzed extensively in the past and meta-analyses are available^2,37^. Therefore, we do not perform a worldwide meta-analysis here but rely on results of these previous work. More in detail, to assess the carbon footprint of meat and dairy products, we gather data from various sources^2,37–50^. Our methodology for retaining data closely follows the protocol of Ref. 37. Requirements to retain sources are that they are issued either through peer-reviewed meta-analysis data or data gathered by recognized national or international agencies. We require that any of these sources contain sufficient information on the methodologies. These are Life Cycle Analysis (LCA) averaged over a national scale or meta-analysis of LCAs that concern worldwide distributed plants/farms.

The LCAs retained share the same functional units (1kg of edible meat or dairy product, 1kg of Fat and Protein Corrected Milk for milk). Where scarce data was available, *e*.*g*. for veal meat, a single study was available for 1 kg of carcass weight, and a 1 : 0.695 conversion ratio to edible weight was used coherently with the study of Ref. 37.

The boundaries of the LCAs retained for this study are from cradle to farm-gate^38,42,44–50^, or beyond. To be more specific they extend to Regional Distribution Centre^37^, to retail^2,39^ or to grave^40^). Transport and other life cycle stages beyond the farm-gate stage for the products considered here (meat and dairy) represent only a small fraction of GHG emissions contributing to the carbon footprint, compared to those emitted from cradle to farm-gate^51^ (in median only 77 *gCO*_2_ *eq*.*/*100*g* edible^37^). For products such as fresh vegetables, these life cycle stages are more relevant^51^. In mode detail, median calculations from cradle to farm-gate^38^ show for a few products slightly *more* important climate change impacts than more complete assessments from *e*.*g*. cradle to Regional Distribution Centre^37^ – potentially due to particular methodological differences in LCA assessment, that are beyond the scope of our work. The additional (relatively) small contributions beyond farm-gate stage for meat and dairy therefore lie within the uncertainty range of the data. In an effort to assess the carbon footprint of the most possible products, we conserve all data with boundaries at least from cradle to farm-gate.

Minimum *C*_m,min_ and maximum *C*_m,max_ median values are taken from world averaged, meta-analysis^2,37^ or national or international agencies^38,39^. Extreme (*C*_min_ and *C*_max_) values are taken from the reported extreme values of aforementioned references or from other national studies^40–45^; or – when data is scarce, for different milk origins (goat, sheep, buffalo), extreme values may be taken from LCAs of several farms^46–50^ (if they exceed other extreme values).

To assess that data was sufficient to investigate differences between products, we used a one way ANOVA test and found a p-value smaller than 0.04 (*<* 10^*−*3^ comparing the 12 main products investigated, 0.035 comparing the 4 milk origins), guaranteeing the validity of the data.

More information on the methods and the data retained in this study can be found in Appendix B.

#### 3. Carbon footprint per g of protein

To assess the carbon footprint per g of protein we divide values of carbon footprint (per g of edible weight) by protein content (per g of edible weight).As both carbon footprint (*C*_min_ − *C*_*m*,min_ − *C*_*m*,max_ − *C*_max_) and protein content are ranges (*P*_min_ − *P*_max_), we can obtain up to 8 ratios.

We present the data with center values as geometric averages based on the carbon footprint medians

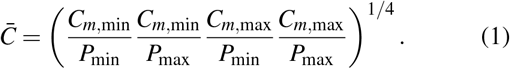

Additionally we report uncertainty ranges as 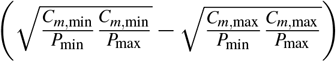. Finally we use extreme values for error bars on plotted displays 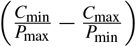. The use of geometric averages is favored over arithmetic averages for better data acknowledgement and to avoid data distortion by extreme values^52–54^. This is coherent with the idea of retaining median values for carbon footprints versus averages. Indeed they give a better representation of the *typical* emissions related to the consumption of a particular food. Specifically, (*i*) we are averaging ratios and the geometric mean treats the numerator and denominator equally. (*ii*) The uncertainty for all the values are rather high (most probably higher than 10%). With an arithmetic mean, a fixed percentage error (say 10%) made on the maximum values would be amplified much more than on the minimum values as they often have different orders of magnitude. (*iii*) Finally, our data is likely to be skewed in some way. Even with all the meta-analyses considered the probability is high that for the same product one might find a sample with higher (resp. lower) values (be it carbon intensity or protein content) than the maximum (resp. minimum) values presented here.

More information on the methods and the data retained in this study can be found in Appendix C.

### B. Special products

We also investigate on a case by case basis the carbon footprint per g of protein of different dairy products. For this detailed investigation, we perform our own meta-analysis. Our analysis protocol follows closely that of Ref. 37. The sources requirements are similar as discussed above, in brief, data sources are issued from LCAs disclosed in peer-reviewed journals with sufficient information on the methods. The LCAs retained share the same functional units. The boundaries of the LCAs are at least from cradle to farm-gate. Most importantly, as this is a close-up investigation, the reference reporting on the carbon footprint should contain the protein content of the cheese or dairy product (whey powder, yogurt) investigated. Exceptions are made for specific cheeses, where the protein content of the product is not directly found in the reference reporting its carbon footprint, but as the cheese is branded or very well identified, its protein content is constant across the market and may be found on sellers databases.

More information on the methods and the data retained in this study can be found in Appendix D.

### C. Regional averages

We also perform analysis of carbon footprint per g of protein on a regional scale. The regions retained for this study are regions were data was most accessible Asia (South and East Asia), Europe, North America, South America and Oceania. A subset of 8 products is studied corresponding to the most consumed products (pork, chicken, beef, lamb, milk, cheese, yogurt and eggs).

#### 1. Carbon footprint

For this investigation, we perform our own meta-analysis. Our analysis relies for the most part on the data base accessible from Ref. 37. Additional (recent, year > 2016) data sources are added to the database in a protocol following closely that of Ref. 37. Additional data sources are issued from LCAs disclosed in peer-reviewed journals with sufficient information on the methods. The LCAs retained share the same functional units. The boundaries of the LCAs are at least from cradle to farm-gate. The full list of data sources is accessible as a Supplementary file.

For each region and each product, median values of carbon footprint 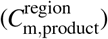 are calculated. One way ANOVA tests were performed for the different products in each region and found a p-value smaller than 0.04 (*<* 10^*−*3^ except for Asia), validating the statistical significance of the different figures between the products. For a few regions and a few products, data availability is too scarce to retain a number. These products correspond to dairy products (cheese or yogurt) in regions where consumption of these products is extremely low relative to other products (Yogurt for Asia, Oceania and South America; Cheese for Asia). As a consequence the exact value of 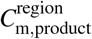 for dietary studies will not affect the results much. In any case, where such data is not available, we evaluate the median carbon footprint of yogurt and cheese based on the carbon footprint for milk in that region, as milk production is the dominant contribution to dairy products carbon footprints^55,56^. In detail, for a specific region *r* we take 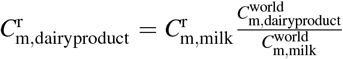 where 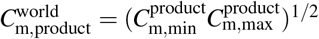, where are the min, max median carbon footprints of a specific product based on world data, as discussed above.

#### 2. Reference diets

We start from reference diets of countries present in the regions investigated. We retain the biggest countries in the regions of interest, namely: China and India (in Asia), the E.U.(in Europe), the United States of America (U.S.A. in North America), Brazil (in South America) and Australia (in Oceania). Average product consumption in each country (or group of countries) is obtained through national or international databases and reports^57–67^. Note that product consumption does not correspond to products *actually* ingested by consumers but are *overestimated*. These numbers do not include food losses at the final stages of the food chain^9^. However, they do correspond to the food that was actually needed for consumption and therefore are the correct amounts to calculate carbon footprint on.

From the reference diets we calculate the total amount of meat and dairy protein intake per diet as

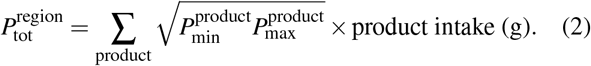

#### 3. Alternative diets exploration

We also explore alternative diets. Our rule of work is to keep the total intake of animal protein 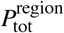 constant across diets. Furthermore we only allow for food items within the initial categories. Our rules are designed to mimic easy swaps for a consumer choosing between food items, and minimal change of diet overall. Within these rules, we investigate 3 alternative diets: (1) a vegetarian diet consisting of dairy and eggs only (ovo-lacto-vegetarian, abbreviated thereafter “Vegetarian”), (2) a low carbon diet containing products that have a low carbon footprint with respect to protein intake, (this choice of products will appear natural after results exploration) namely chicken, yogurt and eggs, termed “Low CO_2_” henceforth, (3) and finally the diet with potentially the lowest possible carbon footprint, containing only chicken (which we will show is the product with the lowest carbon footprint per g of protein), termed “Chicken”.

The amount of the different food items for each specific diet was adjusted such that the relative amounts of the food items are consistent with the relative amounts in the reference diet. Once again, this rule is designed to investigate alternative diets that are as close as possible to actual diets. Accordingly, for each food item, consumption has to be multiplied by a diet factor to meet the goal of conserved total protein intake. For example, for E.U. in the vegetarian diet, the diet factor is 2.5, meaning that an individual would have to ingest 2.5 times more dairy and eggs than average and cut out all meat sources to keep the total amount of protein constant.

#### 4. Carbon footprint per diet

We calculate the carbon footprint per diet using either regional data 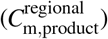 or world data 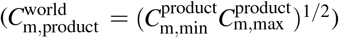. The total carbon footprint is simply

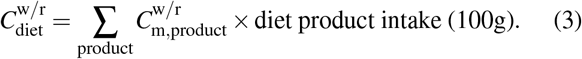

More details on consumption and carbon footprint calculation for each countries (or group of countries) is reported in Appendix F.

## III. MAIN RESULTS: THE CARBON FOOTPRINT PER G OF MEAT AND DAIRY PROTEINS

Figure 3-c and Table I recapitulate the main results of our analysis, showing carbon footprint per g of protein for most common, protein rich, meat and dairy products. The data is sorted from the product with the highest carbon footprint per g of protein to that with the lowest. The results we find are comparable to Refs. 2, 30, and 31 for the few categories investigated in these studies.

**FIG. 3.**
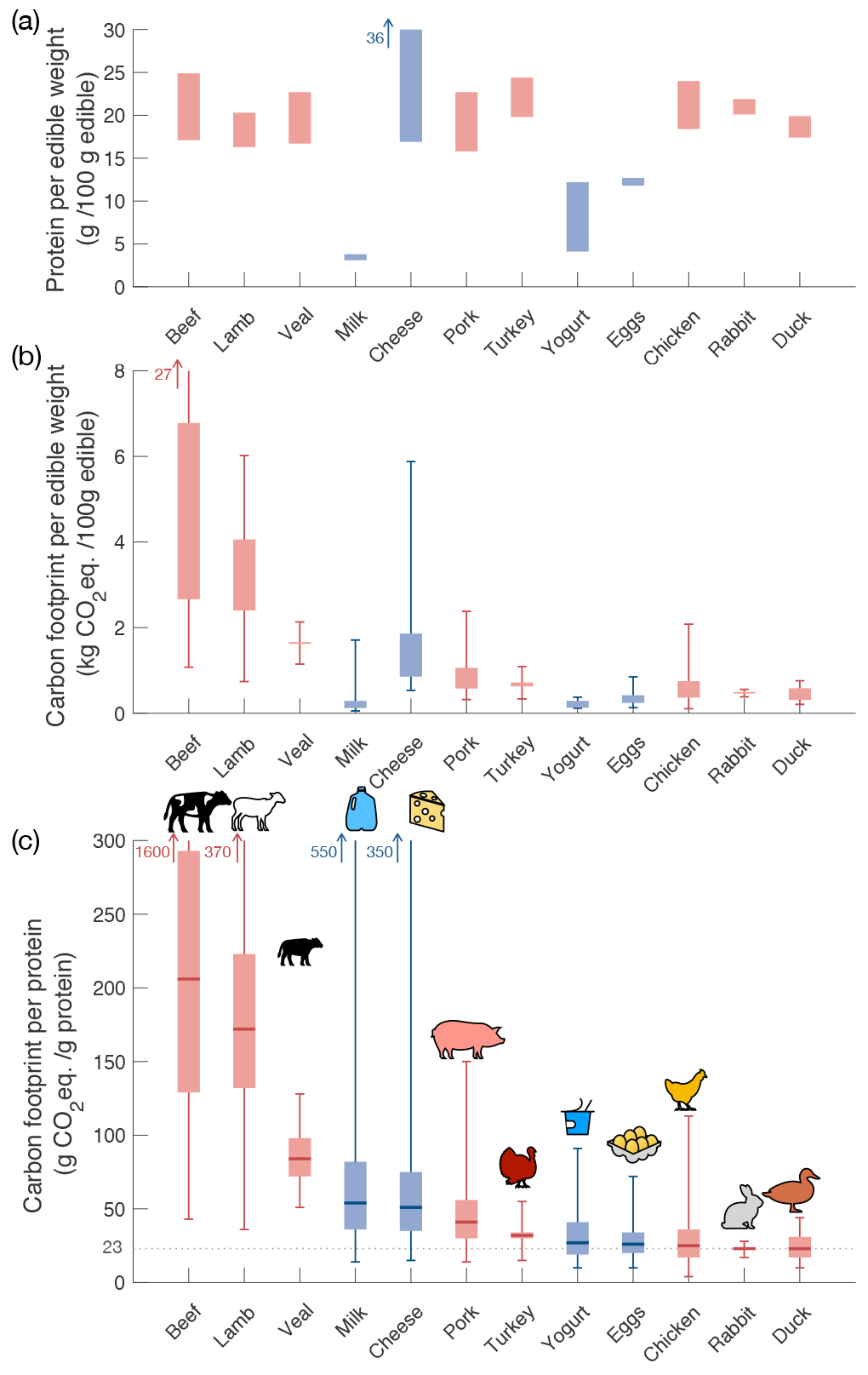
Carbon footprint of animal proteins. (a) Protein content range (b) Carbon footprint range and errorbars and (c) carbon foot-print per g of protein retained and calculated in this study for different meat and dairy products. The dashed grey line in (c) is an indicator line corresponding to the lowest value of carbon footprint per g of protein found in this study.

**TABLE I.** Protein based carbon intensity of common meat and dairy products (sorted from the most impactful to the less). *C* refers to carbon footprint, *P* to protein content and *m* to median. All quantities are given in 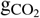 eq./g protein. As detailed in methods, the average carbon intensity 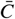 is calculated from the geometric average of the 4 previous columns, while the uncertainty range is given by the geometric average of the 2 first and 2 last columns.

| Product | $\frac{C_{m,min}}{P_{max}}$ | $\frac{C_{m,min}}{P_{min}}$ | $\frac{C_{m,max}}{P_{max}}$ | $\frac{C_{m,max}}{P_{min}}$ | Average $\bar{C}$ |
| --- | --- | --- | --- | --- | --- |
| Beef | 107 | 156 | 272 | 396 | 206 (129-329) |
| Lamb | 118 | 147 | 200 | 249 | 172 (132-223) |
| Veal | 72 | 98 | NA | NA | 84 (72-98) |
| Milk | 32 | 39 | 74 | 90 | 54 (36-82) |
| Cheese | 24 | 51 | 51 | 110 | 51 (35-75) |
| Pork | 25 | 36 | 47 | 67 | 41 (30-56) |
| Turkey | 28 | 32 | 32 | 36 | 32 (30-34) |
| Yogurt | 11 | 32 | 24 | 70 | 27 (19-41) |
| Eggs | 19 | 20 | 33 | 36 | 26 (20-34) |
| Chicken | 15 | 20 | 31 | 41 | 25 (17-36) |
| Rabbit | 21 | 23 | 22 | 24 | 23 (22-24) |
| Duck | 16 | 18 | 29 | 34 | 23 (17-31) |

### A. Insight from carbon footprint per gram of (meat or dairy) protein

#### 1. Protein content range and necessity for a carbon footprint per g of protein

The protein content of the different products is presented in Fig. 3-a. While for most (unprocessed) meats the protein content range is roughly similar, around 20*g/*100 g(edible), the protein content of dairy products is very broad, ranging from 3*g/*100 g(edible) for milk to 36*g/*100 g(edible) for some specific hard cooked cheeses. Within a single food category, the range itself can be very broad (from 17−36*g/*100 g(edible) for cheese products – excluding for now cream cheese or cottage cheese that go as low as 8*g/*100 g(edible)). This highlights the importance of a quantification of carbon footprint per g of protein.

#### 2. Carbon footprint per g of edible food

The carbon footprint per edible weight of the different products is presented in Fig. 3-b. Some food categories have exceptionally large carbon footprints, such as meat from beef, lamb and veal, ranging in average from 2 − 6*kg* CO_2_ eq.*/*100 g(edible)). Other foods have specifically small carbon footprints, in particular lightly processed dairy products such as milk and yogurt, with about 100 − 300*g* CO_2_ eq.*/*100 g(edible). Yet, as mentioned earlier, these products have clearly different protein content as well, and therefore these extreme differences will be greatly reduced when investigated the carbon footprint per g of protein.

#### 3. Carbon footprint per g of protein

In Figure 3-c we observe that some foods that have comparable protein content (such as meats) have very different carbon footprints per g of protein. This is mostly due to the very different carbon footprints per g of edible food of these meats – spanning 2 orders of magnitude from 20*g*CO_2,*eq*_*/g* for chicken, duck and rabbit to 200*g*CO_2,*eq*_*/g* protein for beef. The difference between different meats is mostly due to the fact that some animals (beef, sheep, veal) are *ruminants* and emit large quantities of greenhouse gas through manure emissions. This is not the case for other animals such as pig, chicken, rabbit and duck. Another interesting result is that in general larger animals have a larger carbon footprint per g of protein. A consumer’s oriented take-away rule (in line with low carbon footprint goals) is thus to favor meat from smaller animals. This is also in line with studies demonstrating that the carbon footprint of meat from beef calves consistently increases with slaughter age^44,45^, due to proportionally higher feed intake required at later stages of the animals life.

Figure 3-c is especially useful to compare foods that have very different protein content such as milk and beef. Milk has the lowest carbon footprint per g of edible food among all the foods considered here. Yet it also has the lowest protein content, making direct comparison with meat difficult. Fig. 3-c clearly shows that milk has a relatively high footprint of 54*g*CO_2,*eq*_*/g* protein, with extreme values ranging higher than the average value for beef. This clearly shows that to compare the carbon footprint of protein-rich foods, it is extremely useful to use such methodology. Interestingly, cheese carbon footprint per g of protein ranks very closely to milk, with an impact twice as high as *e*.*g*. chicken. This hints that lacto-ovo-vegetarian diets (abbreviated thereafter to vegetarian), based on high intake of dairy products such as cheese or milk, may not be as effective in reducing carbon footprint as other more carefully designed alternative diets. Such alternative “low carbon diets” could *e*.*g*. include chicken and exclude carbon intensive meats such as beef.

Beyond the products investigated here, there are a number of other meat and other dairy products available on the market. Among these, game meat – often a locally bought meat − may appear as a low carbon alternative. In fact, game meat is not taken into account in national carbon assessments, because the Kyoto protocol considers that game meat is part of the ecosystem and does not contribute to *anthropogenic* carbon emissions^41,68^. Be that as it may, it is interesting to note that ruminants such as deer emit comparable, high amounts of greenhouse gas, much like their mass-produced counterparts, such as beef and lamb^69^.^7^ The variety of dairy products on the market is also quite large and we turn to investigate these in details in the following section.

### B. Specializing into dairy products: milk, cheese, yogurt, whey … and butter

Compared to meat, the variety of dairy products (high in protein content) is quite large: from milk with different skimming contents, to yogurts with added fruit or reduced fat, and the never ending array of cheese options. Furthermore, dairy can be derived from different animal milks. Among all these high protein dairy products, a consumer may wonder which one to choose to achieve the lowest climate change impact. This is what we address in the following paragraphs.

#### 1. Cow or goat milk ?

Milk production around the world originates from different sources. For example, although cheese production across the world is essentially made of cow milk (94%) a small fraction of cheese is made from sheep (3%), goat (2%) and buffalo (1%)^70^. Therefore, one may wonder which source of milk is less carbon intensive among these different animals. Here we review the carbon footprint per gram of protein of these different milks.

Comparing different milk sources is especially interesting since the protein content of milk varies among species – see Fig. 4-a. In particular, sheep milk has a protein content about 2 times larger than cow, goat or buffalo milk. However, the carbon footprint per edible weight of sheep milk production is also the largest among these species – see Fig. 4-b. Overall this results in only slight differences between species when comparing the carbon footprint per g of protein – see Fig. 4-c and Table II. Cow’s milk is the less carbon intensive per g of protein (potentially due to a generally more optimized production line, cow’s being the species most commonly used), closely followed by sheep and goat milk. Finally buffalo milk appears to be the most carbon intensive per g of protein, nearly twice as high in average as cow’s milk. This final comparison comes with some uncertainty as limited data is available for buffalo’s milk.

**TABLE II.** Protein based carbon intensity of various milks (sorted from the least impactful to the most). *C* refers to carbon footprint and *P* to protein content and *m* to median. All quantities are given in 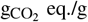 protein. The average carbon intensity is calculated from the geometric average of the 4 previous columns, while the uncertainty range is given by the geometric average of the 2 first and 2 last columns.

| Product | $\frac{C_{m,\min}}{P_{\max}}$ | $\frac{C_{m,\min}}{P_{\min}}$ | $\frac{C_{m,\max}}{P_{\max}}$ | $\frac{C_{m,\max}}{P_{\min}}$ | Average $\bar{C}$ |
| --- | --- | --- | --- | --- | --- |
| Cow's | 32 | 39 | 74 | 90 | 54 (36-82) |
| Sheep's | 20 | 43 | 81 | 171 | 59 (29-118) |
| Goat's | 34 | 42 | 137 | 168 | 76 (36-152) |
| Buffalo's | 89 | NA | NA | NA | 89 (NA) |

**FIG. 4.**
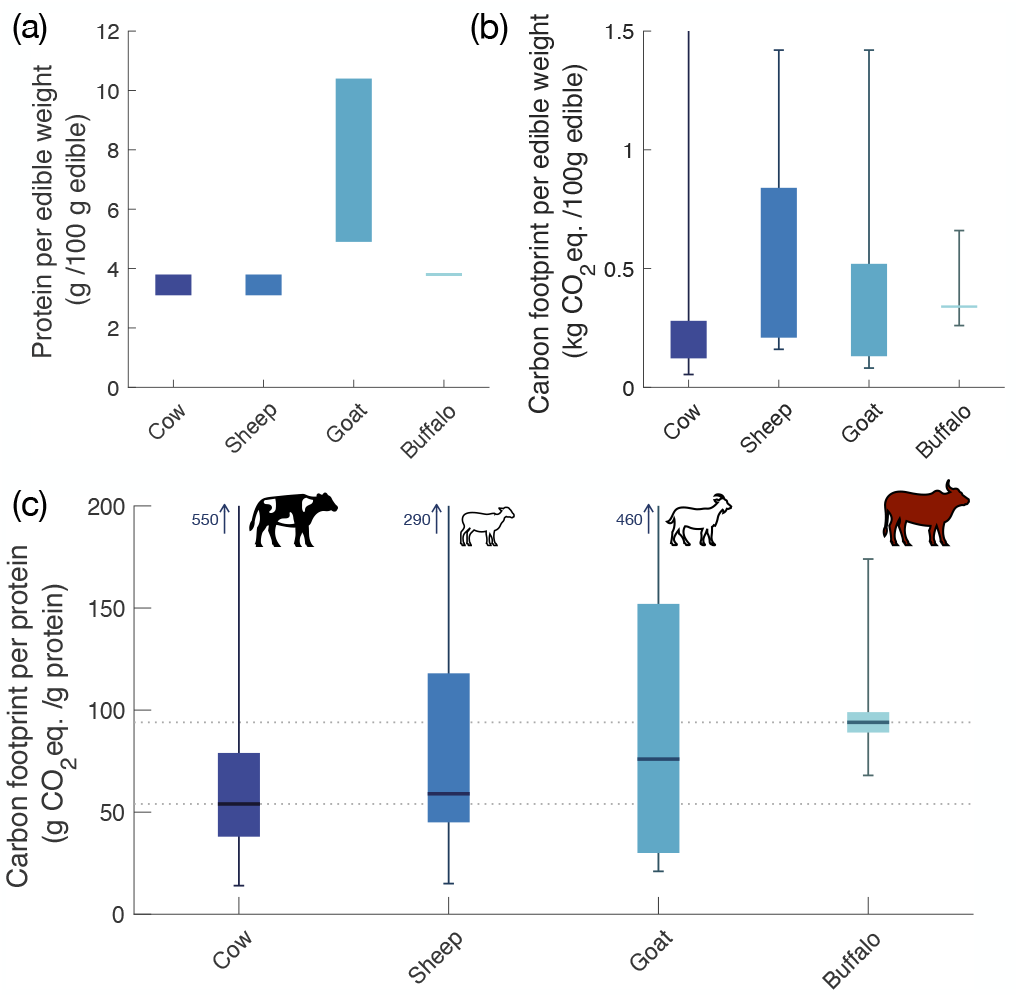
Carbon footprint of various milks. (a) Protein content range (b) carbon footprint range and errorbars and (c) carbon footprint per g of protein retained and calculated in this study for different milks. The dotted lines in (c) are guides for the eye at the average cow and buffalo carbon footprints.

In summary, milks from different species have a comparable carbon footprint per g of protein, with cow’s milk being a little less intensive. Comparing milk from different species is only at its early stage. For example, LCA analysis of milk depends on a correction factor accounting for the typical quality of milk, referred to as the FPCM factor (fat and protein corrected milk). This factor corrects for milk quality between different farms – for example a farm may produce cow milk with a slightly higher protein ratio than another. It is well calibrated for cow and sheep but still under study for goat milk^50^.

Farm by farm analysis giving directly the carbon footprint per g of protein of the milk produced could be done to mitigate this issue, but the protein content of the produced milk is generally not reported.

#### 2. Different cheeses don’t just taste different

Milk is the main primary component of dairy products, and dairy products are extremely varied, especially for cheeses. Cheeses range from fresh cheese to hard cooked cheese, and all possible intermediate compositions. Because cheese preparations are so broad, the protein content of cheeses covers the broadest range of values: from 3− 5*g* protein/100g for fresh yogurt, a few 10*g*/100g for cream cheeses, common cheeses such as cheddar or mozzarella range between 15 − 25*g*/100g and finally aged, very hard cheeses, such as parmesan, hit up to 36*g/*100*g* – see Fig. 5 and Table XI. However, cheeses with a higher protein content generally require more aging and thus have a larger carbon footprint^71^. It is thus natural to wonder whether the added carbon footprint is compensated by the higher protein content. To answer this question we investigate the carbon footprint per g of protein for cheese.

**FIG. 5.**
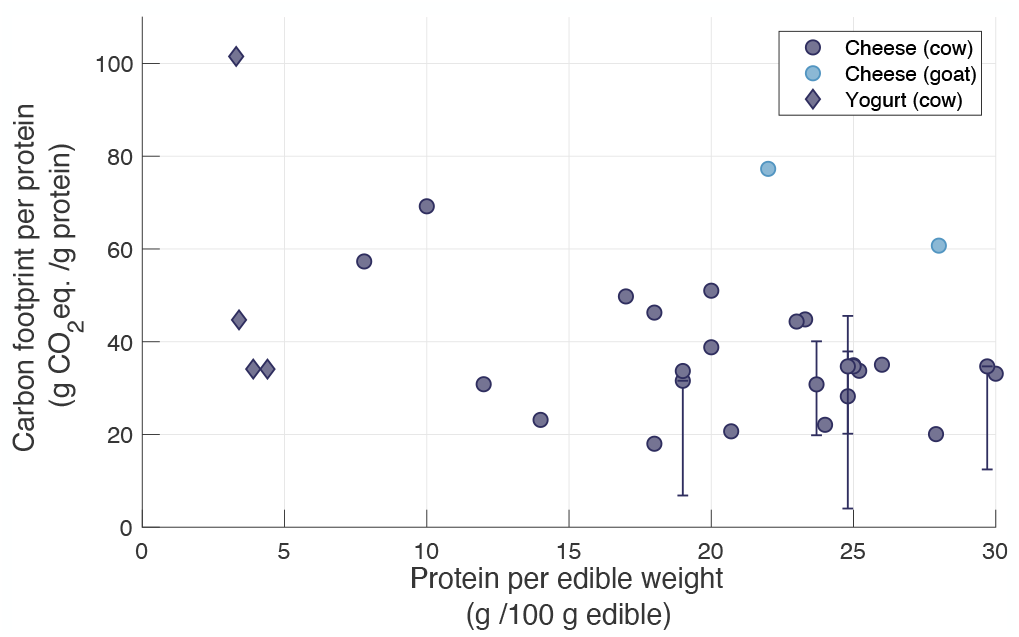
Carbon footprint per g of protein of various dairy products, with a focus on cheese. The data is presented with respect to the protein content of the different dairy products. When confidence intervals were provided by the references investigated, they are reported on the graph.

Raw milk production is the main component of a cheese’s carbon footprint thus most carbon quantification efforts for cheese are focused on reducing the carbon footprint of milk produced for dairy plants^55,56^. In particular, the carbon footprint of cheese strongly depends on whether raw milk was produced on site, or transported – in its liquid or dehydrated state^55^. The second most significant contributor to the carbon footprint of cheese is processing^71^. Interestingly, industrial versus traditional techniques seem to perform quite as well carbon wise^72^. The aging part of processing is the most relevant part^55^. For example, Dalla et al.^73^ compare the carbon footprint of aging for two cheeses, conducting a life cycle analysis restricted to the aging process. They find that through aging protein content increases from 24 to 28g protein/100 g (edible) and carbon footprints rise from 1.32 to 1.61 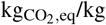 (giving 5.5 to 5.7 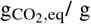 protein). This hints to the fact that additional carbon footprints may well be compensated by higher protein content.

This compensation effect is far from trivial. In fact, one could expect the carbon footprint of proteins from cheese to simply increase concurrently with cheese aging. Yet cheese aging generates a number of co-products (whey, cream, butter, buttermilk *etc*.) to which part of the GHG emissions are also allocated^71,74^. The carbon content of cheese (specifically aged cheese) is significantly dependent on what carbon weight is attributed to those co-products^74,75^. Even in the same plant, differentiating the carbon footprint of two cheeses is quite subtle^71^.

To investigate statistically whether higher protein content compensates for the GHG emissions related to aging, we present data from a number of LCA – see Table XI. We cover a wide range of cheeses, with a wide range of protein contents, and compare their carbon footprint per g of protein in Fig. 5. The correlation coefficient of the data is 0.68 and a least squares linear regression yields a regression coefficient *r*^2^ = 0.47. This confirms that the carbon footprint per g of protein of cheese does not depend significantly on the cheese’s protein content. Therefore, in a consumer’s low carbon perspective, choosing between different cheeses (as a protein source) is not relevant.

#### 3. Whey Powders for protein supplements

The dairy product with the largest protein content is whey protein concentrate, with 80-90g protein/100g^76^. For these products, data availability is extremely limited. Nonetheless, an extensive study allows to establish that whey concentrates emit 0.96 - 1.0 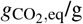 protein^76^ (with LCA boundaries from cradle to farm-gate), see Table. XII. Concentrated whey is therefore one of the animal proteins with the lowest carbon footprint. This is consistent with another study that shows that whey, per protein serving, has one of the smallest carbon footprints among different high protein options^77^ (with similar LCA boundaries).

In contrast, standard whey products (not concentrated) – used for infant formula for instance – have similar carbon footprint per g of protein as cheeses^71,76,78^ (similarly, with LCA boundaries from cradle to farm-gate), see Table. XII.

#### 4. The carbon footprint of butter

Analyzing different varieties of cheese highlights the critical role of co-products of the dairy industry in carbon footprint assessment. Some of these co-products are particularly concentrated, not in protein, but in fat, such as butter (and other creams and oily preparations). Worldwide consumption of these products can not be disregarded. For example, in 2014, the worldwide average butter consumption was 700g/capita/year^70^ (reaching much higher and much lower values in specific countries). With a carbon footprint of 11.52 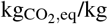 in average^37^, this makes up about 8 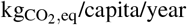 for butter consumption. To put this number in perspective, with 8 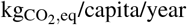, one could alternatively get 11-22 servings of chicken or 1-3 servings of beef (100g steaks, see Table VII).

Interestingly, butter is one of the most carbon intensive sources of fat per kg^2,37^. Therefore, butter may very well be the high-fat product with the highest carbon footprint per g of fat. This sets the question of what are the best products (carbon wise) to obtain fat? An analysis similar to our analysis on proteins, this time comparing products with respect to their carbon footprint per g of fat, could be done to answer this question – yet is beyond the scope of the current study.

## IV. CASE STUDY: CARBON FOOTPRINT OF DIFFERENT DIETS CONTAINING MEAT AND DAIRY

We now turn to investigate how dietary choices may affect the carbon footprint of an individual. In fact, when discussing carbon footprint of the food supply chain, improvements in crop and breeding techniques can have notable impact on the carbon footprint but seem to be insufficient to achieve IPCC targets^3,4,79^. Dietary changes have to be considered to meet this goal. There are many potential dietary choices and numerous authors have investigated the potential positive impact of alternative diets on carbon emissions^2,3,5,79–89^. A detailed investigation of different dietary choices and their carbon footprint is beyond the scope of this study. Instead, we keep a focus on animal proteins from meat and dairy, and investigate *among* these food categories, the carbon footprint of specific choices. First, we consider alternative diets starting from a reference diet (that of the average European) – see Sec. IV A. Then we explore how these dietary choices are more or less effective on carbon footprint reduction starting from different reference diets across the world – see Sec. IV B.

### A. Example of dietary changes based on the European average diet

We start by investigating in detail the carbon footprint of specific dietary choices on a representative diet, the average European Union (EU) diet. The carbon footprint per g of edible product in Europe is reported in Table III. Table IV recapitulates meat and dairy consumption in average in Europe and the content and carbon footprint of different alternative diets. We base our calculations on the data and methodology presented in the Methods Sec. II C.

**TABLE III.** Regional carbon footprint data for products retained in this study.

| Regional carbon intensity for meat or dairy products ($C_m(C_{\min} - C_{\max})$, gCO<sub>2</sub> eq./100g edible) | | | | | |
| --- | --- | --- | --- | --- | --- |
| Meat or dairy Product | Asia (AS) | Europe (E.U.) | North America (NA) | South America (SA) | Oceania (OC) |
| Pork | 720 | 545 | 610 | 120 | 765 |
| Chicken | 740 | 360 | 307 | 373 | 385 |
| Beef | 3526 | 2576 | 2682 | 3392 | 2267 |
| Lamb | 3939 | 2856 | 3268 | 3788 | 1759 |
| Milk | 200 | 127 | 116 | 149 | 114 |
| Cheese | 1351<sup>a</sup> | 880 | 872 | 1445 | 910 |
| Yogurt | 208<sup>a</sup> | 135 | 154 | 155<sup>a</sup> | 119<sup>a</sup> |
| Eggs | 339 | 319 | 446 | 380 | 145 |

<sup>a</sup> computed from world average figures ( $\mu_{\text{world}}^{\text{milk}}, \frac{\mu_{\text{region}}^{\text{milk}}}{\mu_{\text{world}}^{\text{milk}}}$ )

**TABLE IV.**
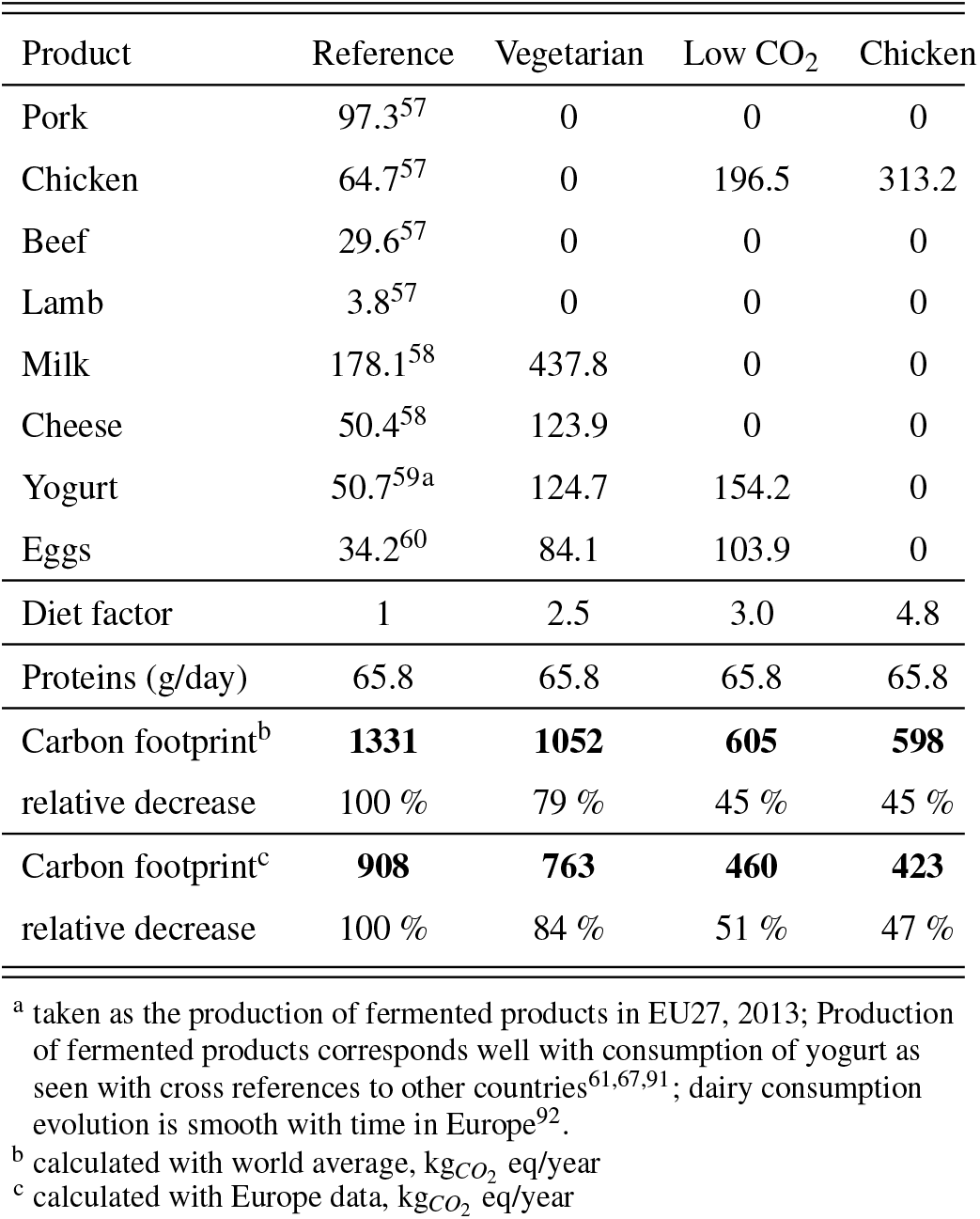
Carbon footprint and protein intake from meat and dairy consumption for a reference **European** diet, and resulting carbon footprint for alternative diets keeping the same total number of proteins from meat and dairy. The product consumptions are all given in g/person/day. The vegetarian diet corresponds here to an ovo-lacto-vegetarian diet. The low CO_2_ diet contains chicken, eggs and yogurt.

The total protein intake coming from meat and dairy in the E.U.is 62.8 g*/*person*/*day. The carbon footprint of the diet based on carbon footprints calculated with world averages is 1331 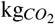 eq/year, while it is lower with regional averages, 908 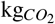 eq/year. This corresponds to overall efficient agricultural techniques in Europe.

The intake of the various food items in the different diets (vegetarian, low CO_2_ and chicken) and the corresponding carbon footprint of the different diets is reported in Table IV. Comparative carbon footprints and diet distribution among products is represented in Fig. 6. Comparing diets based on world averages carbon footprints or on regional data yields very consistent results. We observe that the carbon footprint of the vegetarian diet is only 20% lower than that of the reference diet. This is due to the fact that the vegetarian diet still heavily relies on dairy products. Dairy originates from ruminants and is thus quite impactful as far as carbon footprint is concerned. Comparatively, the low CO_2_ diet achieves a 50% reduction in the carbon footprint. This is interesting because it highlights that – within the rules defined in this study – a vegetarian diet may not be quite as effective as other diets including meat to reduce carbon footprint. Note that here, our low CO_2_ diet includes chicken, but other kinds of meat originating from small animals (duck, rabbit) could work. The chicken only diet achieves a marginal improvement compared to the low CO_2_ diet as the low CO_2_ diet is already quite abundant in chicken.

**FIG. 6.**
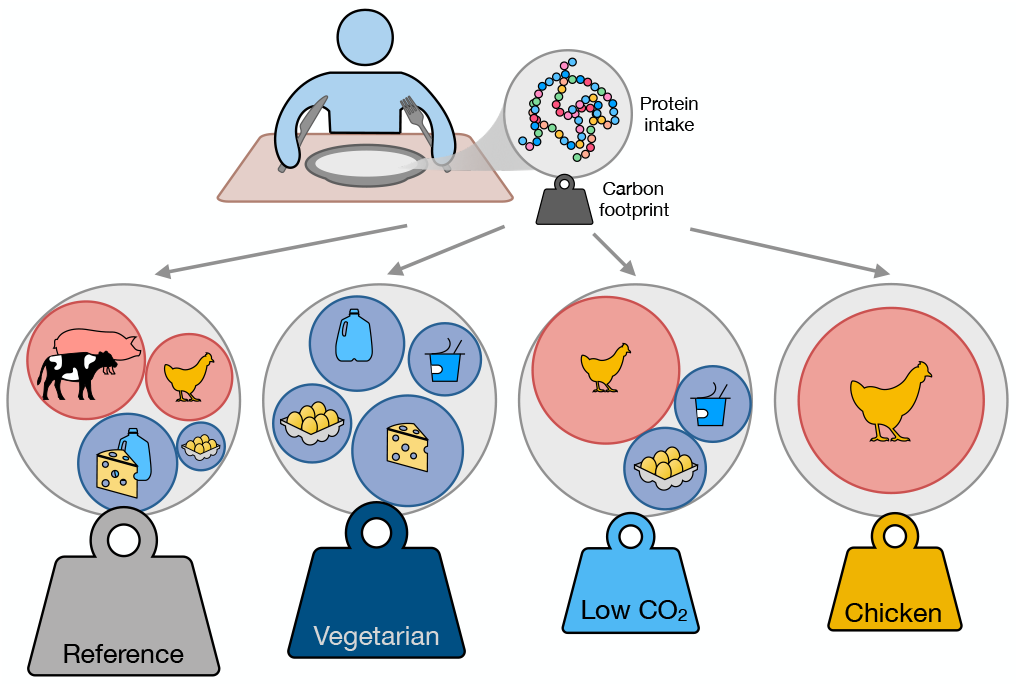
Carbon footprint of different dietary choices starting from a reference E.U. diet. The plates show 4 different diets (reference, vegetarian (with eggs and dairy only), low CO_2_ (with chicken, eggs and yogurt) and chicken). The disks in the plates represent proportional contributions of the various animal proteins to the diets. The carbon weights attached to each plate also have areas proportional to the relative carbon footprints.

The improvement in carbon footprint of diets when shifting from meat and dairy products to poultry was also noted by other works investigating complete or partial diet alternatives^4,81,83,85,88,90^. Dairy rich diets, or diets replacing meat by dairy products are in general not found to yield significant improvement of the carbon footprint of the diet^81,83^. Comparing products solely based on their protein content fails however to take into account the benefits of specific micronutrients that can be found in these products^82^. This could slightly shift the balance, and we discuss these facts in more detail in Sec. V B.

### B. Impact of specific dietary changes across the world

Next, we explore how the efficiency of these alternative diets translates for representative populations across the world. This is quite relevant since carbon footprint reduction when switching diets is dependent on location^4^. Here, we investigate dietary changes for populations in Brazil, in the U.S.A., in Australia, in China and in India. The exact same methodology as for the European diet was applied for these different countries, using both world average and regional carbon footprints – see Table. III. The results of carbon footprints across diets and countries are reported in detail in Appendix E and presented synthetically in Fig. 7.

**FIG. 7.**
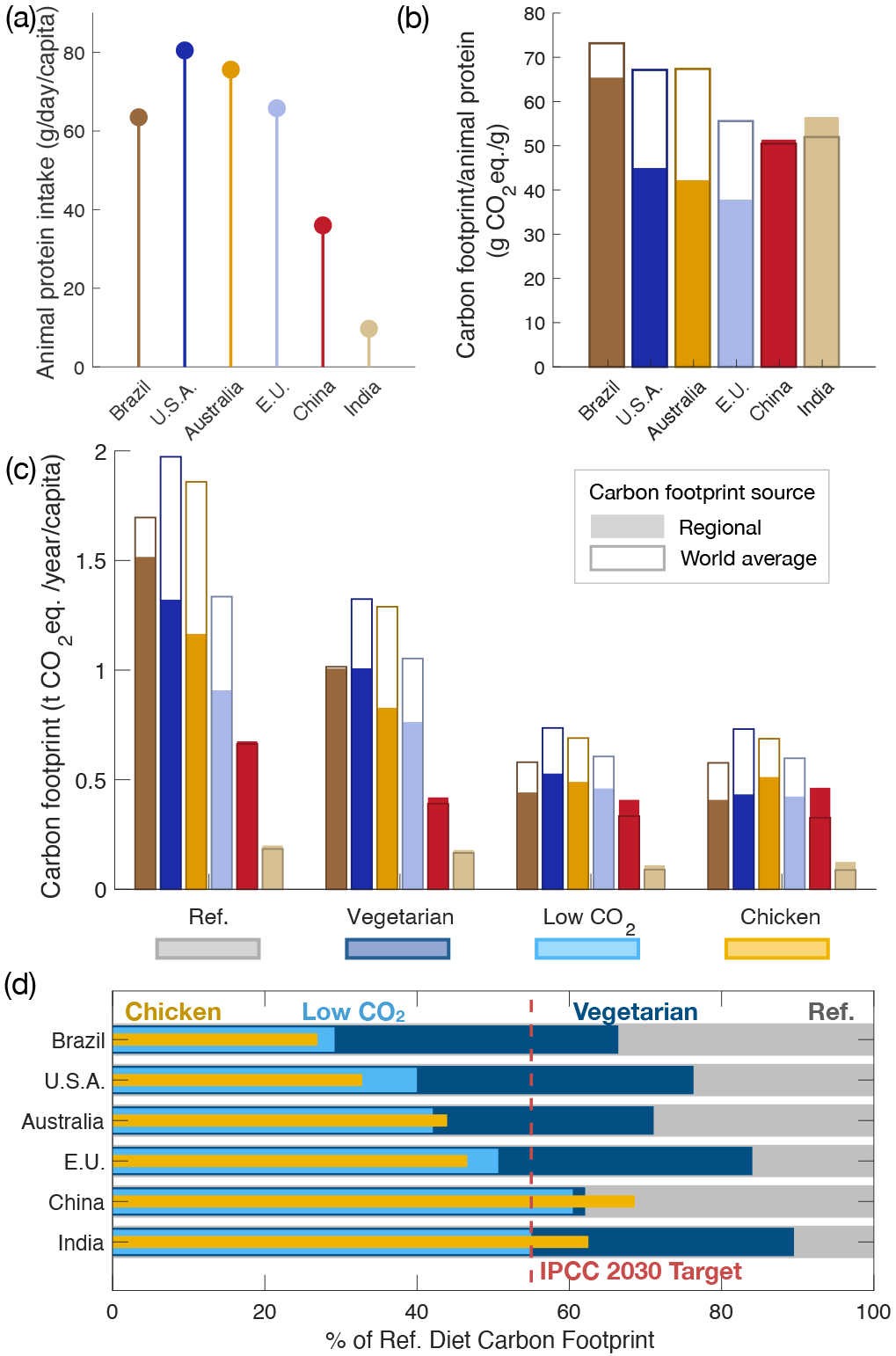
Carbon footprint reduction for different dietary choices in different countries. (a) Daily animal protein intake (from meat and dairy, excluding complementary protein intake in the form of *e*.*g*. protein powder) for 6 representative countries; (b) Average carbon footprint per animal protein; (c) Yearly carbon footprint of the 4 different diets investigated in Fig. 6 for the 6 countries; (d) Relative carbon footprint reduction for the different diets investigated for the 5 countries comparing data calculated with regional carbon footprints.

The choice of countries is purposely done to illustrate the diversity of dietary behaviors. For example, in average, in these countries the protein intake from meat and dairy is very disparate (see Fig. 7-a), ranging from 80 g/day/capita in the U.S.A. to barely 10 g/day/capita in India. Even including 30 % typical losses at the consumer level, this amounts to typically 56 g/day/capita of protein ingested in the U.S.A. which is quite significant compared to dietary recommendations (48 − 64 g/day recommended for a 60 − 80 kg individual − see Sec. I B.). For India, in fact this represents only a small fraction of dietary recommendations.

The average carbon footprint per g of protein for these different countries is also quite different, see Fig. 7-b. The world average data (open boxes in Fig. 7-b) measures how diet composition influences the carbon footprint per g of protein. We find that diets that are rich in meat and especially in beef (such as the Brazilian, the American and the Australian diets) achieve the highest carbon footprint per g of protein. In comparison, diets with quite high amounts of chicken or vegetarian diets (such as Chinese and Indian diets) achieve slightly lower carbon footprints per g of protein (about 30% lower). Investigating the regional carbon footprint per g of protein of the diet (full boxes in Fig. 7-b) shows that more economically developed countries typically achieve better footprints than their world counterparts, mostly due to improved agricultural methods and yields^93^. Eventually the carbon footprint per g of protein of the American and Australian diets (high in beef, but with high yields) slightly outperforms the Chinese, Indian and Brazilian ones (10 − 30%).

Yet to compare diets for climate change mitigation, it is relevant to investigate the overall carbon footprint of the diet and not just the carbon footprint per g of protein. Economically highly developed countries are also those consuming the most meat and dairy, and therefore those that achieve the highest carbon footprint per year per capita, even taking into account regional agricultural efficiency (see Fig. 7-c, reference diet). We will now explore the results of potential dietary changes, keeping as a rule regional data for carbon footprints.

Interestingly, the switch to a vegetarian diet (including dairy and eggs) results in a small carbon footprint reduction (15-30% reduction) and is insufficient in all countries to reach the IPCC 2030 target. It is the most effective (achieving about 30 % carbon footprint reduction) starting from the Brazilian and Chinese reference diets – see Fig. 7-c and d, dark blue. Indeed, beef is a predominant component of the Brazilian diet. Therefore any alternative diet without beef achieves much better than the reference diet. For the Chinese diet, the analysis is different. The vegetarian Chinese diet contains quite low amounts of dairy but high amounts of eggs, because dairy is not a major part of the reference diet. Eggs are quite low in carbon footprint per g of protein compared to dairy products. The vegetarian Chinese diet therefore resembles the “Low CO_2_” diet. Comparatively, the switch to a vegetarian diet in India is quite ineffective as the initial average diet is already nearly vegetarian.

Therefore, it is only natural to seek an alternative diet to the vegetarian diet to see if it is possible to achieve better carbon footprint reductions with alternative choices. We propose a “low CO_2_” diet, that takes into account the results of carbon footprint per g of protein. Worldwide data (Table. I and Fig. 3) suggests the following interesting products for carbon footprint reduction: meat from small animals (poultry, rabbit), eggs and yogurt. We therefore propose a “low CO_2_” diet with eggs, chicken and yogurt only. We find that such dietary change is quite effective for all countries – see Fig. 7-d, light blue – allowing carbon footprint reductions from 40 up to 70 %. These reductions meet the requirements of the IPCC 2030 target. Low CO_2_ dietary changes are especially effective for Brazil, America and Australia as the initial consumption of beef in these countries is relatively high compared to other countries (so the potential reduction is higher).

Finally we explore how the switch to a chicken only diet performs. Chicken is, regarding worldwide averages, the product with the lowest carbon footprint per g of protein, and therefore a natural choice. We find that such a restriction does not improve carbon footprint reduction for all countries – see Fig. 7-d, yellow. For some countries a marginal reduction as compared to the low CO_2_ is achieved, while for others a higher carbon footprint than the “low CO2” diet, or even the vegetarian diet, is reached. In fact, although chicken is, regarding worldwide averages, the product with the lowest carbon footprint per g of protein, it is not necessarily the case in all countries. For example, in China, eggs perform much better than chicken, potentially due to different agricultural management techniques compared to other countries^93,94^. The mix of products in the low CO_2_ diet thus avoids small regional disparities. Overall this demonstrates that a shift to the low CO_2_ diet (*i*) has a drastic impact over the carbon footprint of an individual from meat and dairy proteins and (*ii*) is a consistently good alternative diet throughout large regions of the world.

## V. DISCUSSION: A LOW CARBON FOOTPRINT CONSUMER GUIDE

Building on our efforts to compare protein-rich foods, we now summarize and discuss how our results and methodology can be extended to provide an actual low carbon footprint consumer guide between protein-rich foods.

### A. Low carbon footprint dietary guide

#### 1. The general low CO_2_ diet: chicken, eggs, and yogurt

We have found that dietary choices within meat and dairy that achieve a low carbon footprint overall, reaching the IPCC 2030 Target, include meats from small animals (chicken, duck, rabbit), eggs, and dairy products with little preparation and high protein content (yogurts). Such dietary choices allow for significant carbon footprint reduction in most regions of the world (the Americas, Europe, South and South East Asia, and Oceania). On other parts of the world similar conclusions may likely hold as well but data on carbon footprints of dairy products is insufficient to explore the question.

Yet consumers may be faced with more detailed questions than just what type of meat and dairy to eat but also in *what* quantity and from *what origin*. These are questions we discuss below.

#### Online tool to guide low carbon footprint dietary choices

To guide low carbon footprint choices, beyond generic choices, quantity matters. For example, the consumption of a 125g steak of beef per year may not significantly change the diet of an individual, but 125g a week could. To facilitate science informed decisions, and as part of a dissemination effort, we provide a simple online tool – see Fig. 8 and Ref. 95 – allowing anyone to estimate their carbon footprint from meat and dairy products. The tool allows the user to enter their weekly consumption of the most common meat and dairy products in a single online interface and returns the yearly carbon footprint and daily protein intake. Further details on the online tool implementation and source may be found in Appendix F.

**FIG. 8.**
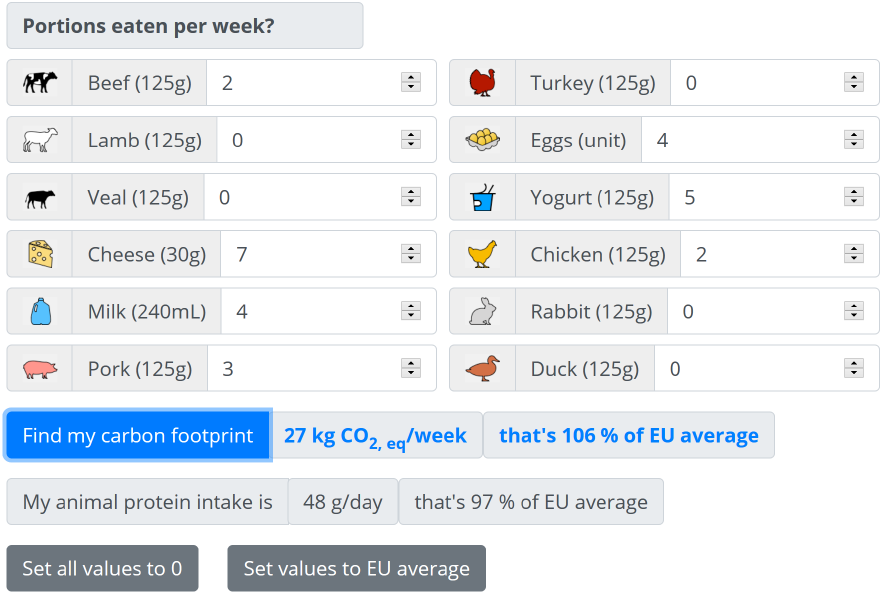
Example use of our online tool^95^ (accessible at http://www.sciriousgecko.com/ArticleMeat.html) for a quick assessment of the carbon footprint of meat and dairy proteins.

#### 2. Local or imported meat ?

Consumers are generally keen on buying and consuming “locally” produced foods^96,97^. Key driving factors include – but are not restricted to – associating health and quality with local products^96,97^, concerns of helping the local economy to thrive and engaging in sustainability^97,98^. Carbon footprint being one of the aspects of sustainability, it is a natural question to ask, when buying meat or dairy, if “local” makes a difference in terms of carbon footprint.

For *ruminant* meat and dairy, compared to the life cycle stages from cradle to farm gate, transport typically represents an infinitesimal fraction of the carbon footprint^2,41,51,99,100^. In fact breeding, crop growing for feeding and manure emissions at the farm represent significantly much more emissions^2,41,100^. As a canonical example, a study showed how dairy (resp. lamb) imported to the United Kingdom (UK) from New Zealand could actually be 2 (resp. 4) times less carbon intensive as dairy (resp. lamb) directly produced in the UK^99, 8^. In this example, the system boundaries are from cradle to farm gate, but includes transport from New Zealand to the UK for the New Zealand meat. The impact of food miles from New Zealand to the UK is greatly compensated by a more efficient production system in New Zealand. In fact the majority of food miles are achieved via refrigerated sea transport, which is largely less intensive than other road or airborne miles^51,99^. As a rule, production methods are the main factor determining *ruminant* meat and dairy proteins’ carbon footprint.

However, when specializing into sub-products of the dairy industry such as cheese, reducing the transport footprint may significantly reduce the carbon footprint of the product overall. In fact, to make cheese, one requires either raw (liquid) milk or curd – a substance obtained from milk after coagulation. Curd is much lighter than the initial total milk required to make it. Therefore transporting curd instead of raw milk before processing can have significant impact on the overall carbon footprint of cheese (15% reduction is reported in^101^ for the production of mozzarella in the Italian dairy sector for an LCA from cradle to grave).

While for ruminant meats and dairy, emissions linked to transport remains under 2% of the total, for poultry and pig meats they average at about 5%^2^. Therefore, consuming locally sourced pork and poultry (or transported with low-carbon footprint means) is consistent with a low-carbon intensity endeavor.

To put in a nutshell – apart from ruminant meats and dairy for which the question has to be sorted on a case by case basis^4,100^ all other protein-rich meat products have generally a lighter carbon footprint if produced locally.

#### 3. Organic or non-organic ?

Consumers also show increased interest in buying organic food products, including for meat and dairy products^102–104^. Similarly, when trying to minimize carbon footprint, one may ask which agricultural method is the best (here we will focus on organic versus non-organic as this is the most available label for a consumer). Comparing different agricultural methods is a challenge due to the limited availability of data and the difficulty to compare different life cycle analysis (LCA) at this level of accuracy. Here we review a few results from authors directly comparing organic and non-organic systems with the same methods.

We first tackle the subject of ruminant meat and dairy. A study on farming in Japan found that the global warming potential of organic versus conventional systems for beef was similar^105^ (for cradle to farm-gate boundaries). In the UK, organic beef and dairy emits about 15% more than conventional farming^41^ (for cradle to farm-gate boundaries), while organic sheep farms emit 42% less CO_2_ equivalent. In Italy, a case study found that organic beef emits even up to 30% more^106^ (for cradle to farm-gate boundaries). A meta-analysis conducted recently reveals that organic cow milk emits 10% less CO_2_ equivalents than conventional^107^ (for cradle to farmgate boundaries). The broad variety of results makes it difficult to conclude on a general trend. Furthermore, when comparing organic versus non-organic ruminant farms, the results strongly depend on the allocation method and also on the method used to account for land use change^108^. They also depend strongly on the specifics of organic farming, and whether modern organic farming techniques are used or not – in particular for manure management^109^.

However the different studies agree on the relative impact of sub-contributions of cow breeding. For instance organic livestock is locally grass-fed with high quality grass (with more clovers and so on)^41,105,107^. Food does not need to be brought from elsewhere, resulting in a decrease of emissions for the organic system. Still, the amount of grass required for grazing is more important, resulting in more land use change; often organic grass is also treated with manure and other organic fertilizers that emit more carbon^41^ – although that depends on manure management^109^. Other authors suggest that the different type of feeding results in more enteric fermentation in the organic feed^106^. Noteworthy, optimization of production by larger farms does not seem to impact significantly the carbon footprint of dairy production^110^ (for cradle to farm gate boundaries). The variability in the relative importance of these factors explains the variability of the results for organic versus non-organic ruminant products.

For non-ruminants such as poultry or pork, data availability is even more scarce. In the Netherlands a study reports that organic pork production emits between 8 and 40% more carbon than conventional^111^, while in the UK organic pork was found to emit 11% less ^41^ (both studies have boundaries from cradle to farm-gate). For poultry in the UK, organic farms emit 46% more CO_2_ equivalents and free-range non organic (versus cage non organic) emit 20% more than conventional^41^. Similarly for eggs in the UK, organic farms emit 27% more CO_2_ equivalents and free-range non organic farms emit 12% more than conventional^41^. In the UK, “optimized” breeding in conventional farms, relying on an efficient use of space, explain the relative better performance of conventional methods^41^. Another important contributing factor is more important grazing in organic systems, that tends to increase emissions^111^.

One important common feature between ruminants and non-ruminants is that the question of the environmental impact of organic versus non-organic agriculture is much broader than just the carbon footprint. Livestock breeding deteriorates soil and water quality (in the form of water and soil eutrophication – increase of nutrient composition, that can disturb the balance of life forms – and acidification). Such deterioration is generally more important in organic farms that rely on more ground use than non-organic farms^41,111^. However non-organic products require in particular more synthetic pesticides^41,109^, which have their own detrimental environmental impact^112,113^. Noteworthy, organic livestock breeding and other “sustainable” breeding approaches – can be beneficial in many more ways (such as introducing nitrogen fixing plants to enhance soil quality), that are further detailed for example in Ref.^109^.

The current data on organic versus non-organic production systems suggests that in general organic meat and dairy production leads to a higher carbon footprint than non-organic, especially via land use change – unless modern techniques are used^109^. Importantly, the “organic” criteria for products strongly depends on respective country laws. Non-organic farms also strive to consider “sustainable” farming approaches that do not necessarily require organic farming^109^. As described above, there is large variability and more in-depth studies are required to assess the climate impact of organic versus non-organic meat and dairy farms.

### B. A nutrition-oriented note on dietary changes

Beyond protein intake, other nutritional aspects should be considered when considering alternative diets^82,86,88^. This is a difficult task, as dietary reference points, *i*.*e*. most current average diets, are not necessarily nutritionally complete^84^. That being said, we still review some of the main nutritional challenges of the diets considered here.

To start with, all the diets investigated, including the reference diets, fail to reach adequate amounts for several nutrients, in particular for iron^114^. Lack of iron is consistently seen in another study investigating micronutrients of a complete average diet^84^. Furthermore, the chicken-only diet – or other alternative diets that do not include dairy – does not provide calcium, coming from dairy in the other diets^82^. This highlights that to achieve a healthy (*i*.*e*. nutritionally complete diet), additional food items should be carefully added to the diet. For dairy-light diets or chicken-only diets, calcium can be found in sufficient amounts with moderate dietary adaptation, for example by consuming more of certain vegetables, fruits or legumes (*e*.*g*. 3 cups of chopped kale bring as much calcium as 1 cup of milk – about 1/3 of the recommended daily allowance)^86,115^. Larger dietary shifts require more careful nutritional adaptations^86^.

The non-reduction of animal protein throughout the alternative diets investigated here is a common downside. Yet, reduction of animal protein intake leads to a number of potential health benefits^85^. For example, the reduction of livestock product consumption by 30 % was projected to decrease the risk of ischaemic heart disease by 15 %^5^. This fact was corroborated by other studies^90^. Moreover, animal protein intake leads to higher blood serum levels of the hormone insulin-like growth factor 1 (IGF-1)^116^. These higher levels are important risk factors in several types of cancer^117,118^ (prostate^119,120^; colorectal^90^, and breast cancer^121^ for example). Furthermore, trading animal proteins for plant-based proteins in a diet comes with an even greater reduction in carbon footprint^2,37^. Plant-based proteins are therefore promising sustainable foods – though comparing their carbon footprint to that of animal proteins is beyond the scope of this study.

All these arguments point to the fact that beyond their content in protein, or in calories, foods should also be compared for their content in micronutrients. For example, the carbon score of dairy could be improved because it does bring important quantities of calcium^82^; similarly pork contains more micronutrients than chicken^84^. Such scoring for diets is still at its early stages and alternative diets – especially vegan diets − should be carefully balanced to fulfill micronutrient targets (Note that a healthy vegan diet reaching all micronutrient targets is possible in developed countries^109,122^ but some studies fail to compare diets where all micronutrient targets are reached^82^). Alternatively, whereas numerous discussions are focused on what micronutrient targets some alternative diets do *not* fulfill; little discussion and scoring is performed on *excessive* micronutrient intake, or potentially long-term disease associated with some foods^122^. For example, although dairy is potentially interesting for its high level in calcium, high dairy intake may be associated with higher risk of prostate cancer^123^ via IGF-1^120^. This is not the case for non-dairy calcium sources. A detailed investigation of micronutrient targets can thus only be performed within entire diet compositions, and with careful set up of scoring measures.

### C. A note on allocation: what about protein allocation ?

When comparing the carbon footprint of different foods, we have noticed how crucial distribution of GHG emissions among sub-products may be (the canonical example of these being distribution between meat and milk in a dairy farm)^2^. In general, when GHG emissions can not be separated, allocation is performed on an economical basis – this is the case in all of the studies investigated here and when data is retained. In a protein-focused perspective, one might expect that eventually milk and beef, two main products generated by cow breeding, should have the same carbon footprint per g of protein. It is therefore only natural to ask how a protein allocation may affect the results investigated here. Performing a complete LCA with protein allocation and comparison with other allocation methods is beyond the scope of this study. Instead, and in line with our pedagogical view, we explore the consequences on a concrete – admittedly very simplified – example.

Let’s consider a dairy farm producing both milk and meat. We assume that 100 (arbitrary) carbon units are necessary for the production of 25 g of meat and 1 kg of milk (typical production ratio^124^). We now seek the carbon footprint per g of protein for milk and meat with two allocation scenarios for the carbon units: (1) protein allocation and (2) economic allocation. Taking data from Table V and VI (≃ 20g protein/100g of edible meat and 3.3 g protein/ 100g of milk) we get 5 g of meat protein per 33 g of milk proteins; a total of 38 g proteins. The protein based allocation (1) gives 100*/*38 = 2.63 carbon units/g of protein be it milk or beef. Taking a price of 0.9$*/kg* of milk and 11$*/kg* for meat^125^ makes 0.9$ of milk and 0.275 $ of meat, a price ratio of 70%. Thus the economic allocation (2) attributes 70 carbon units to milk (respectively 30 to meat), making 70*/*33 = 2.1 carbon units per kg of milk protein and 30*/*5 = 6 carbon units for meat protein. Note that the coarse-grained numbers calculated here give a good representation of more advanced analyses (with boundaries within the dairy system only^124^ or beyond from cradle to retail^2^).

**TABLE V.** Protein Content of common meat-based products. All of the data reported is given without bones^32^, and for raw meat.

| Protein Content of common meat-based products |  |  |
| --- | --- | --- |
| Meat-based product | Protein content range<br>$P_{\min} - P_{\max}$ (g/100g raw) | List of references used |
| Beef | 17.1-24.9 | *Trimmed parts* (lean) 22.5<sup>32</sup>, (fat) 18.9<sup>32</sup> (minced) 19.7<sup>32</sup>, (rumsteak) 20.7<sup>32</sup> (loin) 22.2-24.9<sup>33</sup> (round) 23.4-23.7<sup>33</sup>, (all) 18.4-24.1<sup>10</sup>; *Ribs* 18.8<sup>32</sup>; *Stewing steak* 22.1<sup>32</sup>, 21.2-24<sup>10</sup>; *burgers* 17.1<sup>32</sup>, (lean and not) 17.3-21.9<sup>10</sup> |
| Veal | 16.7-22.7<sup>a</sup> | *Scallops* 20.7<sup>10</sup>, 22.7<sup>32</sup>; *burgers* 16.7-17.2<sup>10</sup>; *other* 18.3-27.3<sup>10</sup>, 16.95-20.92<sup>34</sup> |
| Pork | 15.8-22.7<sup>b</sup> | *Bacon* 16.5<sup>32</sup>, 15.8<sup>32</sup>, 17<sup>10</sup>; *ham* (salami) 18.4<sup>32</sup>, 17.4<sup>32</sup> (ham, 4% fat) 20.9<sup>32</sup> (cooked) 15.1-18.1<sup>33</sup>, 18-21.6<sup>10</sup> (uncured) 24.2-30.4<sup>10</sup>; *trimmed* 18.6-21.8<sup>32</sup> 15.9-22.7<sup>10</sup>; *sausages* 11.9-13.6<sup>32</sup>, 11.8-17.3<sup>10</sup> |
| Chicken | 18.4-24<sup>c</sup> | *Dark meat, such as thighs* 20.9-24.0<sup>32</sup>; *Dark and white* 18.4-23.5<sup>10</sup>; *roasting, meat only* 20.44<sup>34</sup>; *ground* 17.04-17.93<sup>34</sup>; *broilers or fryers, variable content of skin* 17.88- 22.2<sup>34</sup> |
| Turkey | 19.8-24.4<sup>c</sup> | *Dark meat, such as thighs* 24.4<sup>32</sup>, 20.6, 21.28<sup>34</sup>; *White meat, such as breasts and wings* 20.4<sup>32</sup>, 21.28<sup>34</sup>, 20.22<sup>34</sup>; *Dark and white* 19.8-23.4<sup>10</sup>; *Sausage* 18.79<sup>34</sup>; *Ground* 19.66<sup>34</sup>; *other* 15.6-19.7<sup>33</sup>, 18.79<sup>34</sup> |
| Lamb | 16.3-20.3 | *Loin chops, cutlets* 16.3<sup>32</sup>, 17.6<sup>32</sup>; *trimmed, minced* 20.2<sup>32</sup>, 19.1<sup>32</sup>; *shoulder* 17.6<sup>32</sup>, 17.5-20<sup>10</sup>; *other* 20.3<sup>34</sup> |
| Duck | 17.4-19.9 | 19.7<sup>32</sup>, 17.4-19.4<sup>10</sup>, 18.28-19.85<sup>34</sup> |
| Rabbit | 20.1-21.9 | 21.9<sup>32</sup>, 20.4-21.8<sup>10</sup>, 20.05<sup>34</sup> |
| Game meat<sup>d</sup> | 20.7 - 23.7 | *deer, roe, pheasant, boar, rabbit* 20.7-23.7<sup>10</sup>; *bison, deer, boar, rabbit* 21.51, 21.62, 21.79, 22.96<sup>34</sup> |
| Ostrich<sup>d</sup> | 20.2 - 23.7 | 20.2<sup>10</sup> 20.22-23.69<sup>34</sup> |
| Organ meats<sup>d,e</sup> | 7.1 - 21.8 | *Pork* 7.1<sup>32</sup>, 12.1<sup>10</sup> *Beef* 10.3-21.8<sup>10</sup> |

<sup>a</sup> keeping only scallops and burgers to keep only well-identified parts
<sup>b</sup> Removing sausages and ham that differ greatly according to the kind of preparation involved
<sup>c</sup> Removing not well identified parts, and sausages and ground meat that differ greatly in preparation
<sup>d</sup> For these products, little or no data on carbon footprint was found. Especially for game meat, where the footprint is very limited since animals are not tended. We elaborate on game meat in the discussion section.
<sup>e</sup> This corresponds to various organ meats such as liver, tongue, etc.

**TABLE VI.** Protein Content of common dairy products; As an indicative note, a typical egg weighs between 40 and 70 grams^133^, resulting in about 4-9 g of protein per egg.

| Protein Content of common dairy products and eggs |  |  |
| --- | --- | --- |
| Dairy product | Protein content range<br>$P_{\min} - P_{\max}$ (g/100g raw) | List of references used |
| Milk (cow, skimmed to whole) | 3.1-3.8 | 3.3-3.4 (*average, UHT or pasteurized*)<sup>32</sup>, 3.24-3.8<sup>10</sup>, 3.06-4.02<sup>33</sup> |
| Milk (goat) | 3.1-3.8 | 3.22-3.77<sup>10</sup> 3.1 (*average, UHT or pasteurized*)<sup>32</sup>, 3.56<sup>34</sup> |
| Milk (sheep) | 4.9-10.4 | 4.85-10.4<sup>134</sup> 5.4 (*average, raw*)<sup>32</sup>, 5.98<sup>34</sup>, 5.68<sup>10</sup> |
| Milk (Buffalo) | 3.8 | 3.75<sup>34</sup> |
| Fresh cheese (cow) | 7.7-13.3 | 12.6 (*cottage cheese*)<sup>32</sup>, 7.65<sup>10</sup>, 8.55-13.3<sup>33</sup> |
| Soft cheese (cow) | 16.9-25.6 | *Brie* 17.3-22<sup>10</sup> 20.3<sup>32</sup>; *camembert* 21.5<sup>32</sup>, 21<sup>10</sup>; *blue cheeses* 20.5-23.7<sup>32</sup> 19.6<sup>10</sup>; *mozzarella* 18.6<sup>32</sup>, 16.9<sup>10</sup>, 20.9-25.6<sup>33</sup> |
| Soft hard cheese (cow) | 20.4-24.6 | *Reblochon* 20.4<sup>10</sup>; *saint-nectaire* 22.5<sup>10</sup>; *raclette* 24.6<sup>10</sup> |
| Cooked cheese (cow) | 21.5-36.2 | *Cheddar* 24<sup>10</sup>, 25.4<sup>32</sup>, 21.5-25.6<sup>33</sup>; *parmesan* 34.1-34.5<sup>10</sup>, 36.2<sup>32</sup>, 34.1<sup>10</sup>; *comte and other related cooked cheeses* 27.1-28.4<sup>10</sup>; *swiss cheese* 25.7-28.3<sup>33</sup> |
| Fresh cheese (goat, sheep) | 14.8-20.7 | *Feta* 14.8<sup>10</sup> 15.6<sup>32</sup>; *other* 19.8-20.7<sup>10</sup> |
| yogurt (plain to low fat, cow) | 4.1-5.7 | 4.12-4.82<sup>10</sup> 4.8-5.7<sup>32</sup>, 5.25<sup>34</sup> |
| yogurt, greek style (low fat, cow) | 6.9-12.2 | 7.95<sup>10</sup>-9.89<sup>10</sup> 6.89-12.2<sup>33</sup> |
| Eggs (chicken) | 11.8-12.7 | 12.5<sup>32</sup>, 11.8-12.7<sup>33</sup>, 12.7<sup>10</sup> |

We find that milk proteins have similar carbon footprints regardless of the allocation method. In contrast meat proteins have higher carbon footprint with economic allocation, nearly three times higher as for milk proteins. Indeed, in dairy farms, the amount of meat is just so little compared to milk that protein allocation tends to *underestimate* the carbon footprint of meat (and only barely overestimates that of milk). As a result, protein allocation may slightly improve the carbon footprint per g of protein of beef, but not of milk. Therefore, both milk and beef have very high carbon footprints per g of protein (compared to other products such as poultry) regardless of the allocation method. ^9^

In general, allocation by the amount of protein (method 1) or (more commonly used) by the amount of energy (calorie content) is not relevant. For example, in many situations the same initial compound may be used for outputs that are not comparable protein-wise or calorie-wise. For example milk, can be used to make whey protein (very high in protein, quite low in energy) or butter (very low in protein, very high in energy). A protein based-allocation would therefore have butter be nearly carbon-free^126^. Carbon allocation based on the relative price of the products – *economic allocation* (method 2) does not suffer from these limitations. In fact, economic allocation has the advantage of drawing more carbon intensity to more demanded products. It also lightens carbon weights of less demanded co-products such as whey or straw. For further comparison of allocation methods we refer the reader to Ref. 2, 101, 124, 126, and 127.

## VI. CONCLUSION AND OUTLOOK

In summary, we have introduced a methodology to compare the carbon footprint of protein rich foods, in particular of meat and dairy protein. Our results show that ruminant meat and dairy have a high carbon footprint per g of protein; while other meats (such as pig and poultry) and protein-rich, lightly processed, dairy (such as yogurt) have a much lower carbon footprint. Importantly, this methodology and the data generated allows to guide low carbon footprint dietary choices within meat and dairy. Interestingly, a change to ovo-lacto-vegetarian diet results in a low improvement of the carbon footprint. A change to a low CO_2_ diet, containing small poultry, yogurt, and eggs results in a drastic, 50 %, improvement, allowing to reach the IPCC 2030 target^1^. Such an alternative diet is easily achievable for a consumer wishing to maintain its total consumption of meat and dairy proteins, and allows significant improvement.

Furthermore, we have investigated several other consumer oriented questions; such as choosing between local or imported, organic or non-organic, and within the variety of dairy products. These investigations point to a limited data availability, showing that some consumer oriented questions are hard to answer at this stage. Nonetheless, among milk origins we find that cow’s milk has the lowest carbon footprint compared to other ruminants. We also find that cheeses have comparable carbon footprints per g of protein regardless of aging. Finally, we identified that locally sourced poultry, or pork may have a significantly reduced footprint, as well as dairy processed directly on the farm.

To reach IPCC targets, the low carbon diet would not be sufficient since such drastic improvements in food emissions can not necessarily be obtained over all food sources^37^. Alternative food sources, and in particular alternative protein sources (plant-based or from fish), should be investigated and compared in similar ways to offer consumer-friendly perspectives. This is the aim of future work. Furthermore, although our study was focused solely on carbon footprint, meat consumption, and in particular red meat consumption, has a high environmental impact with respect to water, pesticide and fertilizer usage, ocean acidification, toxic emissions in the air and land eutrophication^41,109,112,113,128,129^.

As outlined in the nutritional discussion in Sec. V B, our study investigates solely the carbon footprint with respect to protein content and does not account for other nutritional aspects. Beyond micronutrient targets, and as highlighted by a number of authors, many other factors come into play. For example when comparing protein rich foods it has been noted that not all protein sources are equivalent because some are easier to digest^77,130^. Furthermore, factors such as very local dependencies of carbon footprint (from one region of a country to another)^87^, economic cost of the alternative diet^86,87^ and cultural adequacy^86^ are very relevant points to address when considering alternative diets. These factors require careful introduction of scoring measures, and all participate in understanding how to best mitigate climate change.

## ACKNOWLEDGEMENTS

The authors are indebted to Amaury Hayat and William Legrand for sparking interest in the topic.

## APPENDICES

### Appendix A protein content of meat and dairy retained for this study

#### A.1. Measuring the protein content of meat

After meat has been cleared from bones – sometimes trimmed from fat – the amount of pure nitrogen contained is measured using a number of chemical reactions. A conversion factor is then used to relate the pure nitrogen content and the nitrogen contained originally in proteins in the food (purple circle in the amino acids of Fig. 2)^32^. This allows to quantify the protein content of the food. The conversion factor most widely used today, in particular used to calculate the data we report below, relies on an early study^35^. However, it is to be noted that this conversion factor is an early estimate that does not properly take into account the various nitrogen contents of proteins^131^ and a different factor is strongly recommended by scientists today^132^. To ensure consistency of our study, we will still use data resulting from the older factor, noting that the difference between the two factors is only 20% and does not vary much among the food categories investigated.

#### A.2. Protein content of meat and dairy products investigated in this study

We report here values of proteins found in meat and dairy products from various national databases^10,32–34^. Protein content of common meat-based products may be found in Table V and of dairy products in Table VI. In each table; the protein content range is the minimum to maximum of protein content that we have retained.

### Appendix B: carbon footprint of meat and dairy retained for this study

We report here values of carbon footprint for the production per gram of meat and dairy products from various sources. Carbon footprint per 100 g of edible food of common meat-based products is given in the Appendix B in Table VII and of dairy products in Table VIII. In each table, we highlight the carbon footprint range that we have retained.

**TABLE VII.** Carbon footprint data of meat products retained in this study. The range of carbon footprints for each product is made out of four numbers: the lowest single value, the lower median value, the higher median value, the highest single value found in meta-analyses or systematic reviews.

| Meat-based product | Carbon intensity for meat-based products |  |
| --- | --- | --- |
|  | Carbon footprint range | List of meta-analysis used, with carbon footprint data |
|  | (gCO<sub>2</sub> eq./100g edible) | (gCO<sub>2</sub> eq./100g edible) |
| | [Min – Median<sub>min</sub>–Median<sub>max</sub> – Max]<br>$C_{min} - -C_{m,min} - C_{m,max} - -C_{max}$ | (**bold font** corresponds to retained data for carbon footprint range) |
| Beef | 1074 – 2661-6780 – 26920 | 2700<sup>40</sup> (single LCA, USA, average methods); 3100<sup>42,43</sup> (Average, Canada); **2661 (1074-10950)**<sup>a,37</sup> (meta-analysis); 6040 (3760-**26920**)<sup>b,2</sup> (meta-analysis); 2860 ± 30%<sup>38</sup> (Average, national agency, France); **6780** (1100-11050)<sup>c,39</sup> (world average emissions, FAO); |
| Veal | 1148 – 1640-NA – 2132 | **1280-1650**<sup>44</sup> (different age of slaughter, typical Danish/Swedish farms)<sup>d</sup>; ; 1600-1990<sup>45</sup> (different age of slaughter, typical E.U.farms); **1640 ± 30%**<sup>38</sup> (Average, national agency, France) |
| Pork | 320 – 577-1060 – 2380 | 1210<sup>40</sup> (single LCA, USA, average methods); **577 (320-1186)**<sup>a,37</sup> (meta-analysis); **1060** (690-**2380**)<sup>b,2</sup> (meta-analysis); 589<sup>38</sup> (Average, national agency, France) |
| Lamb | 740 – 2400-4060 – 6020 | 3920<sup>40</sup> (single LCA, USA, average methods); 2558 (1005-5670)<sup>a,37</sup> (meta-analysis); **4060** (2370-**6020**)<sup>b,2</sup> (meta-analysis, no distinction between Lamb and Mutton); 3300± 30%<sup>38</sup> (Average, national agency, France); **2400 (740-4970)**<sup>c,39</sup> (world average emissions, FAO) |
| Chicken | 106 – 365-750 – 2080 | 690<sup>40</sup> (single LCA, USA, average methods); **365 (106-998)**<sup>a,37</sup> (meta-analysis); **750** (400-**2080**)<sup>b,2</sup> (meta-analysis); 475± 30%<sup>38</sup> (Average, national agency, France) |
| Turkey | 334 – 628-717 – 1090 | **1090**<sup>40</sup> (single LCA, USA, average methods); **717 (334-849)**<sup>a,37</sup> (meta-analysis); **628± 30%**<sup>38</sup> (Average, national agency, France) |
| Duck | 207 – 309-583 – 758 | **309 (207-410)**<sup>a,37</sup> (meta-analysis); **583± 30%**<sup>38</sup> (Average, national agency, France) |
| Rabbit | 382 – 470-486 – 558 | **470 (382-558)**<sup>a,37</sup> (meta-analysis); **486± 30%**<sup>38</sup> (Average, national agency, France) |

<sup>a</sup> numbers reported here correspond to: median (min-max) as given in Ref.<sup>37</sup>;
<sup>b</sup> numbers reported here correspond to: median (5th percentile-95th percentile) as given in Ref.<sup>2</sup>. Ref.<sup>2</sup> does not provide minimum or maximum values used and hence we choose to use (5th percentile-95th percentile) as good representatives of these values
<sup>c</sup> numbers reported here correspond to: average (min-max) where the average is the LCA for the total world production, and the variation of footprint corresponds to varying yields across the world. The LCA is from cradle to retail.
<sup>d</sup> Using carcass weight to edible weight values reported in Ref.<sup>37</sup>

**TABLE VIII.** Carbon footprint data for dairy products retained in this study. The range of carbon footprints for each product is made out of four numbers: the lowest single value, the lower median value, the higher median value, the highest single value found in meta-analyses or systematic reviews.

| Dairy product | Carbon intensity for dairy products |  |
| --- | --- | --- |
|  | Carbon footprint range | List of meta-analysis used, with carbon footprint data |
|  | (gCO<sub>2</sub> eq./100g edible) | (gCO<sub>2</sub> eq./100g edible) |
| | [Min – Median<sub>min</sub> - Median<sub>max</sub> – Max]<br>$C_{\min} - C_{m,\min} - C_{m,\max} - C_{\max}$ | (**bold font** corresponds to retained data for carbon footprint range) |
| Cheese (Cow) | 533 – 855-1860 – 5880 | 1347<sup>40</sup> (single LCA, USA, average methods); **855 (533-1635)**<sup>a,37</sup> (meta-analysis); **1860 (1020-5880)**<sup>b,2</sup> (meta-analysis) |
| Yogurt (Cow) | 117 – 131-288 – 374 | 217<sup>40</sup> (single LCA, USA, average methods); **131 (117-200)**<sup>a,37</sup> (meta-analysis); **288 ± 30%**<sup>38</sup> (Average, national agency, France) |
| Milk (Cow) | 54 – 122-280 – 1710 | 129 (**54-750**)<sup>a,37</sup> (systematic review); 270 (150-700)<sup>b,2</sup> (meta-analysis); **122 ± 30%**<sup>38</sup> (Average, national agency, France; 106-123<sup>41</sup> (Table 59, LCA non-organic vs organic, Average, England and Wales); **280 (130-1710)**<sup>c,39</sup> (world average emissions, FAO) |
| Eggs (chicken) | 130 – 241-420 – 850 | 483<sup>40</sup> (single LCA, USA, average methods); 346 (**130-600**)<sup>a,37</sup> (meta-analysis); **420 (290-850)**<sup>b,2</sup> (meta-analysis); **241**<sup>38</sup> (Average, national agency, France); 525-700<sup>41</sup> (Table 58, 100% cage non-organic vs organic, egg weight 50 g, Average, England and Wales); |
| Milk (Sheep) | 160 – 209-840 – 1420 | **840 (160-1420)**<sup>c,39</sup> (world average emissions, FAO); 320 (200-520)<sup>d,46</sup> (12 farms, variation of footprint corresponds to varying yields in different farms, cradle to farm-gate LCA); **209**<sup>38</sup> (Average, national agency, France) |
| Milk (Goat) | 81 – 131-520 – 1420 | 104-140<sup>50</sup> (16 representative farms, cow's milk FPCM correction factor retained to be consistent with other studies, variation of footprint corresponds to varying allocation scenarios, cradle to farm-gate LCA); 267 (112-505)<sup>d,49</sup> (17 farms, variation of footprint corresponds to varying yields in different farms, cradle to farm-gate LCA); **520 (160-1420)**<sup>c,39</sup> (world average emissions, FAO) 89 (**81-103**)<sup>d,48</sup> (5 farms, variation of footprint corresponds to varying yields in different farms, cradle to farm-gate LCA); **131**<sup>38</sup> (Average, national agency, France) |
| Milk (Buffalo) | 260 – 340 - NA – 660 | **340 (260-660)**<sup>c,39</sup> (world average emissions, FAO); 360-375 (287 - 520)<sup>d,47</sup> (6 farms, cradle to farm-gate LCA); |

<sup>a</sup> numbers reported here correspond to: median (min-max) as given in Ref. 37;
<sup>b</sup> numbers reported here correspond to: median (5th percentile-95th percentile) as given in Ref.<sup>2</sup>. Ref.<sup>2</sup> does not provide minimum or maximum values used and hence we choose to use (5th percentile-95th percentile) as good representatives of these values
<sup>c</sup> numbers reported here correspond to: average (min-max) where the average is the LCA for the total world production, and the variation of footprint corresponds to varying yields across the world. The LCA is from cradle to retail.
<sup>d</sup> numbers indicated here correspond to average value (farm min - farm max) as given in adjoining references. When several average values are given they correspond to variability across possible allocation scenarios;

### Appendix C: Calculating carbon footprint per g of protein

We present in Tables. IX and X extreme values of carbon footprint per g of protein as calculated using the extreme retained values of carbon footprint in Appendix B.2.

**TABLE IX.** Protein based carbon intensity: extreme values.

| Meat-based/dairy product | Protein based carbon intensity: extreme values |  |  |
| --- | --- | --- | --- |
| | $C_{\min} / P_{\max}$ | $C_{\max} / P_{\min}$ | Geometric average carbon intensity <sup>a</sup> |
|  | (gCO<sub>2</sub> eq./g protein) | (gCO<sub>2</sub> eq./g protein) | (gCO<sub>2</sub> eq./g protein) |
| Beef | 43 | 1574 | 194 |
| Lamb | 36 | 369 | 172 |
| Veal | 51 | 128 | 84 |
| Milk | 14 | 552 | 54 |
| Cheese | 15 | 348 | 51 |
| Pork | 14 | 150 | 41 |
| Turkey | 15 | 55 | 32 |
| Yogurt | 10 | 91 | 27 |
| Eggs | 10 | 72 | 26 |
| Chicken | 4 | 113 | 25 |
| Rabbit | 17 | 28 | 23 |
| Duck | 10 | 44 | 23 |

<sup>a</sup> Data reported from Table I

**TABLE X.** Protein based carbon intensity for milks: extreme values.

| Milk product | Protein based carbon intensity: extreme values |  |  |
| --- | --- | --- | --- |
| | $C_{\min} / P_{\max}$ | $C_{\max} / P_{\min}$ | Geometric average carbon intensity |
|  | (gCO<sub>2</sub> eq./g protein) | (gCO<sub>2</sub> eq./g protein) | (gCO<sub>2</sub> eq./g protein) <sup>a</sup> |
| Cow's | 14 | 552 | 54 |
| Sheep's | 15 | 290 | 59 |
| Goat's | 21 | 458 | 76 |
| Buffalo's | 68 | 174 | 89 |

<sup>a</sup> Data reported from Table II

### Appendix D: carbon footprint per g of protein of different type of cheeses

We present in Table XI the protein based carbon intensity of different cheeses and in Table XII of more varied dairy products.

**TABLE XI.** Protein based carbon intensity of different cheeses (starting from cow’s milk, unless otherwise mentioned).

| Type of cheese | Protein based carbon intensity of different types of cheeses |  |  | Notes |
| --- | --- | --- | --- | --- |
|  | P<br>(g<br>protein/100g) | C<br>(gCO<sub>2</sub> eq./100g) | C/P<br>(gCO<sub>2</sub><br>eq./g<br>protein) |  |
| Yellow cheese low fat | 30 | 993<sup>135</sup> | 33 |  |
| Grana Padano | 29.7 | 1030 (max with diff. allocations 1690)<sup>136</sup> | 35 (max 57) | hard cooked, dry matter allocation as central value |
| Pecorino artisanal | 28 | 1700<sup>72</sup> | 61 | goat cheese, hard cooked |
| Emmental | 27.9<sup>10</sup> | 560<sup>38</sup> | 20 | hard cooked |
| Yellow cheese | 26 | 911<sup>135</sup> | 35 |  |
| Dutch Cheese | 25.2 | 850<sup>137</sup> | 34 | semi-hard |
| Hushallsost | 25<sup>138</sup> | 873<sup>139</sup> | 35 |  |
| Gouda | 25<sup>34</sup> | 867<sup>140</sup> | 35 |  |
| Cheddar | 24.8 | 700 (range with diff. allocations 460-1300)<sup>78</sup> | 28 (19-52) | Semi-hard, economic allocation as central value |
| Cheddar | 24.8 | 860 (range with diff. allocations 590-1220)<sup>71</sup> | 35 (24-49) | Semi-hard, economic allocation as central value |
| Cheese (generic) | 24 | 530<sup>31</sup> | 22 | canadian meta-analysis |
| Mozzarella | 23.7<sup>33</sup> | 730 (range with diff. allocations 510-990)<sup>71</sup> | 31 (22-42) | semi-hard, protein content may vary significantly |
| San Simon da Costa | 23.3<sup>141</sup> | 1044<sup>142</sup> | 45 |  |
| Casin (hard) artisanal | 23<sup>143</sup> | 1020<sup>127</sup> | 44 |  |
| Pecorino (industrial) | 22 | 1700<sup>72</sup> | 77 | goat cheese, hard cooked |
| Camembert | 20.7<sup>10</sup> | 428<sup>38</sup> | 21 |  |
| Soft cheese | 20 | 776<sup>144</sup> | 39 | meta-analysis |
| Franxon (artisanal) | 20<sup>145</sup> | 1020<sup>127</sup> | 51 | semi-hard |
| Gorgonzola | 19 | 600 (max with diff. allocations 1070)<sup>146</sup> | 32 (max 56) | hard cooked, dry matter allocation as central value |
| Cheese | 19 | 640<sup>76</sup> | 34 | semi-hard to hard, meta-analysis |
| Fresh cheese | 18<sup>144</sup> | 324 | 18 | meta-analysis |
| White cheese | 18 | 833<sup>135</sup> | 46 |  |
| Mould cheese | 17 | 846<sup>135</sup> | 50 |  |
| Cottage cheese | 14 | 324<sup>31</sup> | 18 | canadian meta-analysis |
| Cottage cheese | 12 | 370<sup>135</sup> | 31 |  |
| Cream cheese | 10 | 692<sup>135</sup> | 69 |  |
| Cream cheese low fat | 7.8 | 447<sup>135</sup> | 57 |  |

**TABLE XII.** Protein based carbon intensity of different dairy products (starting from cow’s milk, unless otherwise mentioned). The carbon intensity reference contains a reference of the protein content of the product investigated.

| Type of product | Protein based carbon intensity of different types of dairy products |  |  | Notes |
| --- | --- | --- | --- | --- |
|  | P<br>(g<br>protein/100g) | C<br>(gCO<sub>2</sub> eq./100g) | C/P<br>(gCO<sub>2</sub><br>eq./g<br>protein) |  |
| Yogurt | 4.4 | 150<sup>31</sup> | 34 | canadian meta-analysis |
| Yogurt low fat | 3.9 | 133<sup>135</sup> | 34 |  |
| Yogurt | 3.4 | 152<sup>135</sup> | 45 |  |
| Yogurt | 3.3 | 335<sup>144</sup> | 102 | meta-analysis |
| Whey protein concentrate (special) | 90 | 17360<sup>135</sup> | 193 |  |
| Whey protein concentrate | 80 | 16400<sup>135</sup> | 205 |  |
| Whey powder | 30 | 1010<sup>31</sup> | 34 | canadian meta-analysis |
| Whey powder | 25 | 740<sup>76</sup> | 30 | meta-analysis |

### Appendix E: typical dietary intakes and comparison of carbon footprint of different diets equivalent in protein

We present in Tables XIII-XVII the carbon footprint and protein intake from meat and dairy consumption for reference and alternative diets for various countries.

**TABLE XIII.** Carbon footprint and protein intake from meat and dairy for a reference **American** diet, and alternative diets. Product consumptions are all given in g/person/day.

| Product | Reference | Vegetarian | Low CO<sub>2</sub> | Chicken |
| --- | --- | --- | --- | --- |
| Pork | 63.0<sup>57</sup> | 0 | 0 | 0 |
| Chicken | 136.2<sup>57</sup> | 0 | 315.6 | 382.9 |
| Beef | 71.5<sup>57</sup> | 0 | 0 | 0 |
| Lamb | 1.4<sup>57</sup> | 0 | 0 | 0 |
| Milk | 181.4<sup>61</sup> | 587.7 | 0 | 0 |
| Cheese | 49.7<sup>61</sup> | 161.1 | 0 | 0 |
| Yogurt | 16.6<sup>61</sup> | 53.9 | 38.6 | 0 |
| Eggs | 39.3<sup>62</sup> | 127.4 | 91.2 | 0 |
| Diet factor | 1 | 3.2 | 2.3 | 2.8 |
| Total protein<sup>a</sup> | 80.5 | 80.5 | 80.5 | 80.5 |
| Carbon footprint<sup>b</sup> | **1 973** | **1 324** | **736** | **731** |
| relative decrease | 100 % | 67 % | 37 % | 37 % |
| Carbon footprint<sup>c</sup> | **1 321** | **1 008** | **528** | **433** |
| relative decrease | 100 % | 76 % | 40 % | 33 % |

<sup>a</sup> calculated, g/day<sup>b</sup> calculated with world average, kgCO<sub>2</sub> eq/year<sup>c</sup> calculated with North America data, kgCO<sub>2</sub> eq/yearTABLE XIV. Carbon footprint and protein intake from meat and dairy for a reference **Chinese** diet, and alternative diets. Product consumptions are all given in g/person/day.

**TABLE XIV.** Carbon footprint and protein intake from meat and dairy for a reference **Chinese** diet, and alternative diets. Product consumptions are all given in g/person/day.

| Product | Reference | Vegetarian | Low CO<sub>2</sub> | Chicken |
| --- | --- | --- | --- | --- |
| Pork | 83.3<sup>57</sup> | 0 | 0 | 0 |
| Chicken | 31.8<sup>57</sup> | 0 | 75.8 | 171.3 |
| Beef | 10.4<sup>57</sup> | 0 | 0 | 0 |
| Lamb | 8.5<sup>57</sup> | 0 | 0 | 0 |
| Milk | 39.2<sup>63</sup> | 144.2 | 0 | 0 |
| Cheese | 0.1<sup>63</sup> | 0.2 | 0 | 0 |
| Yogurt | 9.4<sup>63</sup> | 34.6 | 22.4 | 0 |
| Eggs | 62.7<sup>64</sup> | 230.9 | 149.6 | 0 |
| Diet factor | 1 | 3.7 | 2.4 | 5.4 |
| Total protein<sup>a</sup> | 36.0 | 36.0 | 36.0 | 36.0 |
| Carbon footprint<sup>b</sup> | **663** | **391** | **334** | **327** |
| relative decrease | 100 % | 59 % | 50 % | 49 % |
| Carbon footprint<sup>c</sup> | **675** | **419** | **408** | **463** |
| relative decrease | 100 % | 62 % | 60 % | 69 % |

<sup>a</sup> calculated, g/day<sup>b</sup> calculated with world average, kgCO<sub>2</sub> eq/year<sup>c</sup> calculated with Asia data, kgCO<sub>2</sub> eq/yearTABLE XV. Carbon footprint and protein intake from meat and dairy for a reference **Brazilian** diet, and alternative diets. Product consumptions are all given in g/person/day.

**TABLE XV.**
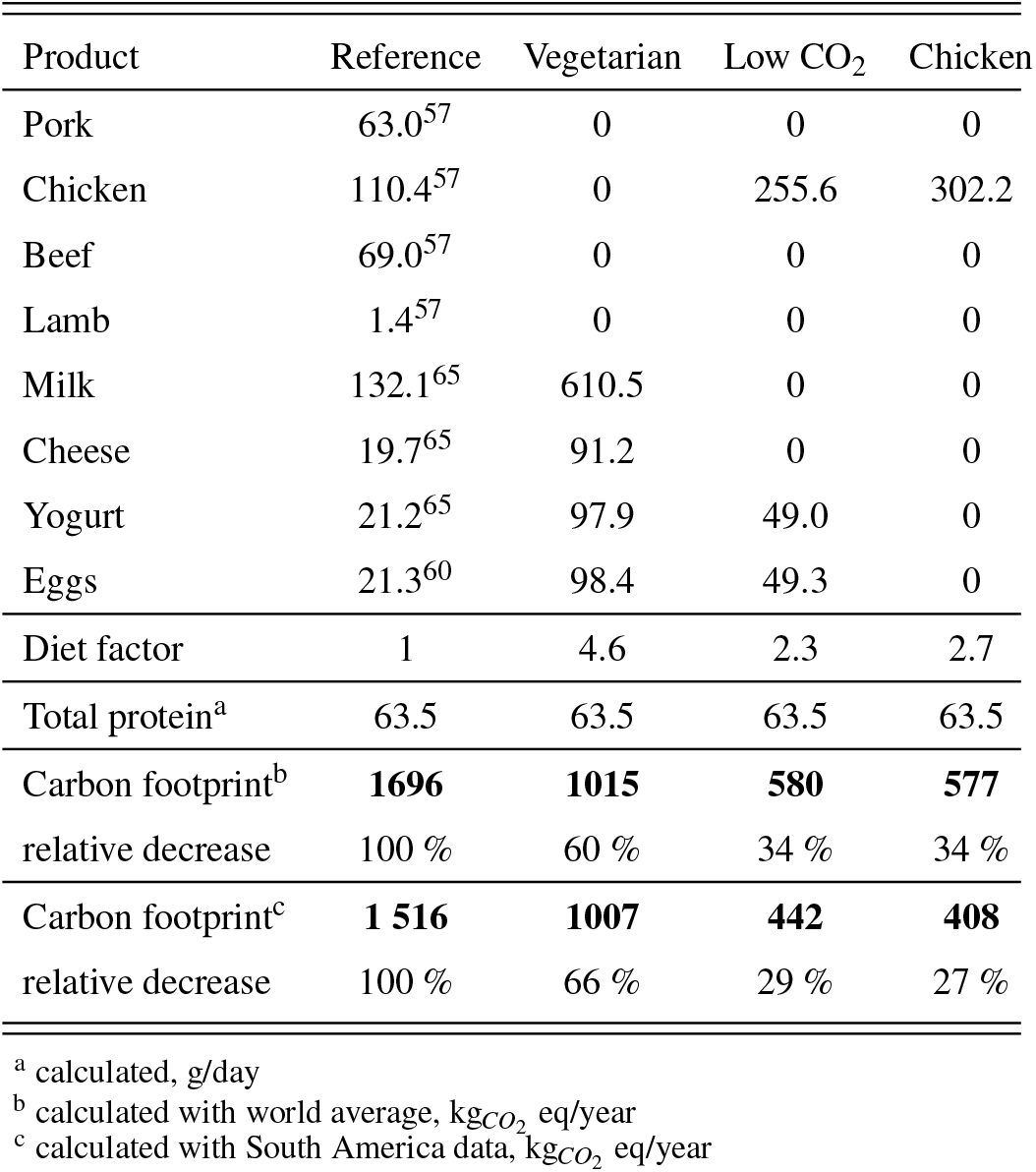
Carbon footprint and protein intake from meat and dairy for a reference **Brazilian** diet, and alternative diets. Product consumptions are all given in g/person/day.

**TABLE XVI.** Carbon footprint and protein intake from meat and dairy for a reference **Australian** diet, and alternative diets. Product consumptions are all given in g/person/day.

| Product | Reference | Vegetarian | Low CO<sub>2</sub> | Chicken |
| --- | --- | --- | --- | --- |
| Pork | 58.6<sup>57</sup> | 0 | 0 | 0 |
| Chicken | 116.2<sup>57</sup> | 0 | 301.4 | 359.5 |
| Beef | 54.8<sup>57</sup> | 0 | 0 | 0 |
| Lamb | 19.2<sup>57</sup> | 0 | 0 | 0 |
| Milk | 275.9<sup>66</sup> | 885.6 | 0 | 0 |
| Cheese | 37.3<sup>66</sup> | 119.6 | 0 | 0 |
| Yogurt | 24.7<sup>66</sup> | 79.2 | 64.0 | 0 |
| Eggs | 22.9<sup>60</sup> | 73.4 | 59.4 | 0 |
| Diet factor | 1 | 3.2 | 2.6 | 3.1 |
| Total protein<sup>a</sup> | 75.6 | 75.6 | 75.6 | 75.6 |
| Carbon footprint<sup>b</sup> | **1 859** | **1 289** | **690** | **687** |
| relative decrease | 100 % | 69 % | 37 % | 37 % |
| Carbon footprint<sup>c</sup> | **1 165** | **828** | **490** | **512** |
| relative decrease | 100 % | 71 % | 42 % | 44 % |

<sup>a</sup> calculated, g/day<sup>b</sup> calculated with world average, kgCO<sub>2</sub> eq/year<sup>c</sup> calculated with Oceania data, kgCO<sub>2</sub> eq/year

**TABLE XVII.** Carbon footprint and protein intake from meat and dairy for a reference **Indian** diet, and alternative diets. Product consumptions are all given in g/person/day.

| Product | Reference | Vegetarian | Low CO<sub>2</sub> | Chicken |
| --- | --- | --- | --- | --- |
| Pork | 0.5<sup>57</sup> | 0 | 0 | 0 |
| Chicken | 6.6<sup>57</sup> | 0 | 21.6 | 46.1 |
| Beef | 1.4<sup>57</sup> | 0 | 0 | 0 |
| Lamb | 1.4<sup>57</sup> | 0 | 0 | 0 |
| Milk | 129.9<sup>65</sup> | 162.9 | 0 | 0 |
| Cheese | 6.6<sup>65</sup> | 8.2 | 0 | 0 |
| Yogurt | 6.3<sup>67</sup> | 7.9 | 20.7 | 0 |
| Eggs | 8.9<sup>60</sup> | 11.2 | 29.2 | 0 |
| Diet factor | 1 | 1.3 | 3.3 | 7.0 |
| Total protein<sup>a</sup> | 9.7 | 9.7 | 9.7 | 9.7 |
| Carbon footprint<sup>b</sup> | **184** | **166** | **90** | **88** |
| relative decrease | 100 % | 90 % | 49 % | 48 % |
| Carbon footprint<sup>c</sup> | **200** | **179** | **110** | **125** |
| relative decrease | 100 % | 90 % | 55 % | 63 % |

<sup>a</sup> calculated, g/day
<sup>b</sup> calculated with world average, kgCO<sub>2</sub> eq/year
<sup>c</sup> calculated with Asia data, kgCO<sub>2</sub> eq/year

### A. Appendix F: an online tool to guide dietary choices

We detail here the methods used to calculate an individuals carbon footprint based on meat and dairy consumption (accessible at http://www.sciriousgecko.com/ArticleMeat.html). Based on input data of weekly consumption, the tool returns the total carbon footprint of the products (based on world averaged carbon footprints 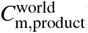), comparing it to the average European Union (E.U.) value (based on world averages) – see Sec. IV. It also gives the corresponding daily protein intake, comparing it to the E.U. average value. Fig. 8 shows an example close to the typical E.U. diet with some modifications.

The data used to compute the carbon footprint and protein intake is taken from Tables V, VI, VII and VIII with a methodology similar to the one detailed in Sec. V. To compare to average E.U. data, we must take into account food losses. In fact, average E.U. consumption data are based on retail sails and not on consumer consumption. We therefore correct the carbon footprint obtained from the user’s consumption by adding a 30% factor, consistently accounting for food losses within the approach by Shepon *et al*.^9^. Note that food losses especially for meat can be much higher (up to 96% for beef). The source code is freely available on the Website^95^. Future releases including geographical origin and type of farming are the object of future work.

## Footnotes

1 food loss at retail stores, in restaurants and household waste which have been estimated to be at least 30% in weight^9^

2 starting from 115 g/day up to 340 g/day gives 230− 680 kcal/day from meat; that is reduced by 30% to account for food loss and finally compared to the typical total intake 2000 kcal/day

3 This is established by several national health agencies. One review notes however that the lowest end of the acceptable macronutrient range (10% of total calories coming from proteins according to the dietary reference intakes^11^) is actually equivalent to about 1.05 g*/*kg*/*day^16^. This is quite larger than 0.8 g*/*kg*/*day

4 Yet a calcium deficient diet *can* impact bone health^18^

5 Other essential nutrients however can not be found in plants, such as Vitamin B12, that is produced by bacteria^26^

6 Similar numbers are found in Europe^28^

7 Keeping for the assessment only manure emissions into consideration: Cattle emit 0.1 kg CH_4_ / kg live weight/ year (taking 53 kg CH_4_ emitted per head per year per cattle, with an average live weight of 550kg^43^). Deer emit a comparable 0.17 kg CH_4_ / kg live weight / year (with 20 kg CH_4_ per head per year per deer, with an average live weight of 120kg).

8 Doing the simple ratio of their numbers 2849*/*688 ≃ 4.1 of the respective carbon intensities for lamb; and 2921*/*1423 ≃ 2 for milk solids.

9 Note that the numbers obtained here show that for economic allocation, beef proteins have a carbon footprint 3 times as high as milk proteins. Yet in Table. VII we find that beef proteins have a carbon footprint 4 times as high is milk proteins. This is due in particular to the fact that beef does not come solely from dairy farms^2,42^.

